# A T6SS DNase effector induces nuclear DNA Damage to trigger apoptosis via activation of the cGAS-STING-TNF axis

**DOI:** 10.1101/2024.09.19.613802

**Authors:** Li Song, Lei Xu, Pengfei Zhang, Shuying Li, Yixin Zhao, Zhenkun Shi, Ruiqi Ma, Yongdong Li, Yao Wang, Gehong Wei, Xihui Shen

## Abstract

In the relentless battle for survival, bacteria have evolved sophisticated weaponry like the Type VI secretion system (T6SS) to not only outcompete rivals but also manipulate host cell processes to their advantage. Apoptosis, a controlled form of programmed cell death without triggering inflammatory responses, is often exploited by pathogens. While many bacterial pathogens are known to induce apoptosis, the role of T6SS effectors in this process remains unexplored. In this study, we discovered that *Yersinia pseudotuberculosis* (*Yptb*) utilizes its T6SS to secrete a DNase effector, TkeA, which induces apoptosis in host cells. Mechanistically, the translocation of TkeA into host cells causes nuclear DNA damage, leading to the release of fragmented DNA into the cytoplasm. This, in turn, activates the DNA-sensing cyclic GMP–AMP synthase (cGAS)/stimulator of interferon genes (STING) pathway. The activation of the cGAS-STING pathway by TkeA subsequently triggers apoptosis in host cells via extrinsic pathways, with tumor necrosis factor (TNF) signaling playing a critical role. Additionally, TkeA also mediates interbacterial competition, promoting the colonization of *Yptb* in mice’s gut. Our findings reveal that the T6SS-secreted DNase effector TkeA acts as a trans-kingdom effector, enhancing interbacterial competition and inducing apoptosis in host cells by activating the cGAS-STING-TNF axis. This discovery adds a new dimension to the mechanisms employed by bacterial pathogens to manipulate host cell death pathways by using their secretion systems.

**GRAPHICAL ABSTRACT:** 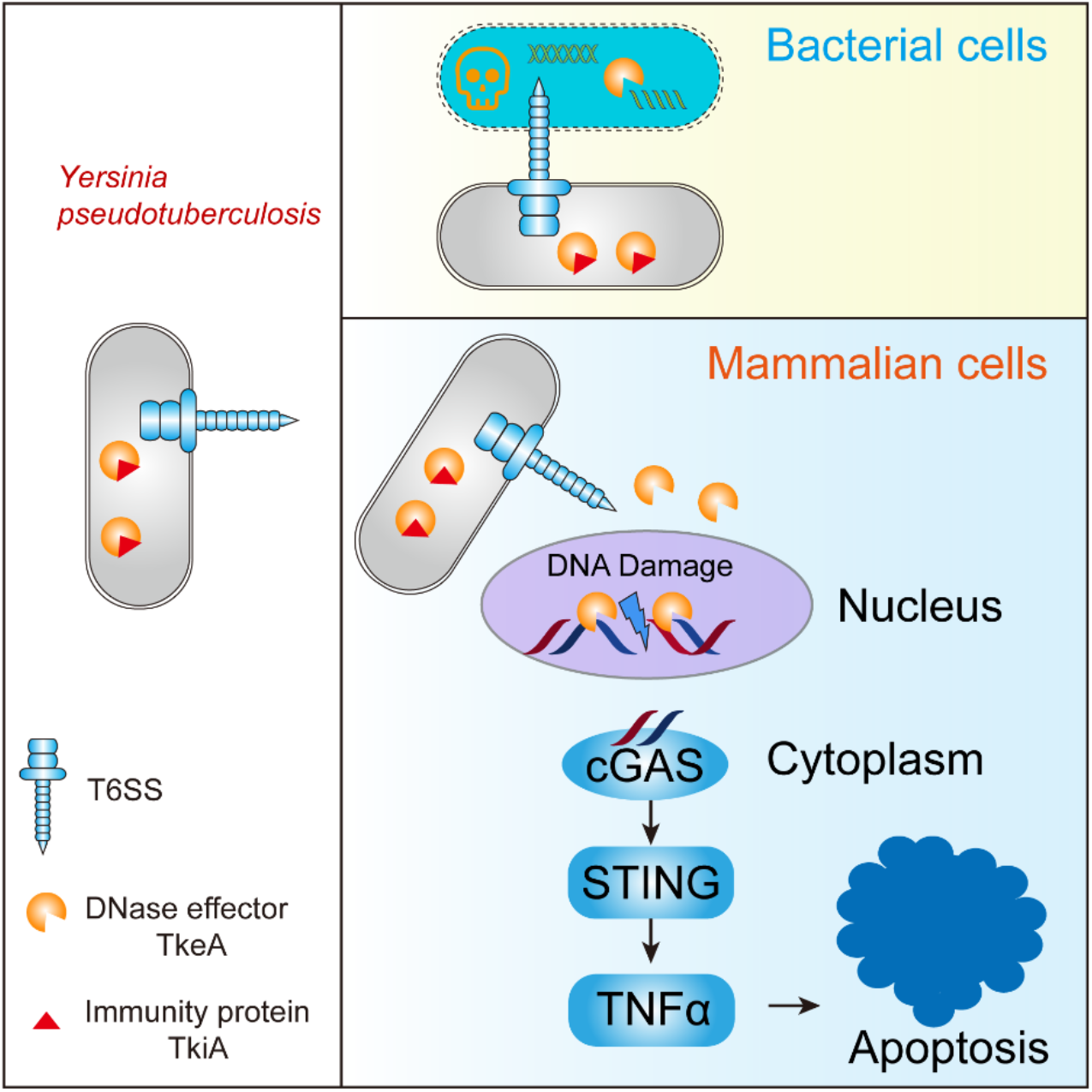

**HIGHLIGHTS:**

• *Yptb* T6SS secretes a trans-kingdom DNase effector TkeA
• TkeA degrades bacterial DNA to mediate interbacterial competition
• TkeA causes nuclear DNA damage in host cells to induce apoptosis
• TkeA induces apoptosis via the cGAS-STING-TNF axis

## INTRODUCTION

Amid the intricate balance of life and death within our cells, apoptosis emerges as a critical guardian, orchestrating the deliberate elimination of infected or damaged cells to maintain tissue homeostasis. This tightly regulated form of programmed cell death (PCD) not only curbs the spread of infection by eradicating compromised cells but also ensures that bacteria cannot exploit these cellular environments for replication.^1^ Unlike other forms of cell death, apoptosis is a non-inflammatory process that can be initiated by both extracellular and intracellular cues. The intrinsic apoptotic pathway is regulated by the Bcl-2 family of proteins, which drive mitochondrial outer membrane permeabilization (MOMP), leading to cytochrome C release and the subsequent activation of CASPASE 9—an event that cascades into the activation of executioner CASPASEs 3/7.^2–4^ External signals, such as tumor necrosis factor alpha (TNFα), can also instigate apoptosis via the extrinsic pathway.^5^ While TNF signaling is primarily known for its role in modulating inflammatory responses through MAPK and NF-κB pathways, it can pivot towards inducing cell death when inflammation is curtailed.^6^ Upon the binding between an external apoptotic TNFα and its corresponding cell membrane receptor TNF receptor (TNFR), signaling complexes are formed and trigger apoptosis by the deubiquitylation and phosphorylation of RIPK1, which includes CASPASE 8 and eventually triggering CASPASE 3 activations.^7^

DNA damage is a potent inducer of apoptosis, often leading to irreversible cell-cycle arrest and p53- dependent intrinsic apoptosis.^8^ DNA damage occurs not only due to normal cellular processes like replication and metabolism but also in response to external factors such as radiation, stress, and microbial infection.^9^ DNA damage-induced apoptosis during bacterial infections is an essential aspect of the host defense strategy.^10,11^ Importantly, the induction of DNA damage is often a strategic mechanism employed by bacteria to manipulate host cellular functions, rather than merely a byproduct of infection.^12^ Bacterial pathogens can induce DNA damage through various mechanisms. Some bacteria secrete toxins that have direct genotoxic effects, such as cytolethal distending toxins (CDTs), colibactin, and indolimines.^13,14,15^ Colibactin and indolimines are small-molecule metabolites,^15,16^ while the CDTs are produced by several Gram-negative bacteria, including *Escherichia coli* and *Campylobacter jejuni*.^17^ CDTs directly induce double-strand breaks (DSBs) in host DNA, resulting in DNA damage that can lead to cell cycle arrest and apoptosis.^14^ These CDT toxins possess a DNase-like activity that cleaves host DNA, thereby initiating a cascade of cellular responses aimed at either repairing the damage or triggering apoptosis if the damage is irreparable.^15–17^ However, how these DNA damage-inducing toxins trigger apoptosis remains largely unknown.

Under normal conditions, DNA is confined within the nucleus and mitochondria. However, during events such as infection, cellular stress, or DNA damage, fragments of DNA can escape into the cytosol, leading to the activation of the DNA sensing pathway. Cyclic GMP-AMP synthase (cGAS) is a universal DNA sensor in mammalian cells that detects cytosol DNA to trigger immune responses by activating the stimulator of interferon genes (STING) protein.^18,19^ This initiates a signaling cascade that leads to the production of type I interferons (IFNs) and other pro-inflammatory cytokines via the phosphorylation of TANK-binding kinase 1 (TBK1) and interferon regulatory factor 3 (IRF3).^18,19^ In addition to its immunomodulatory function, many studies have demonstrated that cGAS-STING signaling also leads to PCD, with apoptosis being extensively studied.^20–25^

The first study indicates that STING-dependent apoptosis involves a BAX-IRF3 complex that promotes intrinsic pathway apoptosis.^20^ Additionally, the cGAS-STING cascade can trigger apoptosis through various mechanisms, including ER stress, NLRP3 inflammasome activation and signaling through NF- κB, IRF3, and IFNs.^26^ Specifically, cGAS-STING-induced ER stress activates the unfolded protein response (UPR) and interacts with the Ca^2+^ sensor STIM1.^27^ Activation of the cGAS-STING pathway also leads to the production of type I interferons (IFNs), which can initiate apoptosis via the JAK-STAT and PI3K-AKT pathways.^28^ Moreover, the cGAS-STING pathway can activate the NF-κB signaling cascade, leading to increased apoptosis by upregulating the p53-upregulated modulator of apoptosis (PUMA).^29^ STING also induces apoptosis by triggering the expression of pro-apoptotic genes.^22,23^ A recent study found that STING interacts with SAM68, leading to IRF3-dependent mitochondrial apoptosis independently of IFNAR signaling.^30^ Since cGAS can recognize leaked self-DNA,^31,32^ the compromise of DNA integrity during bacterial infection may induce apoptosis via the cGAS-STING pathway. However, no studies have specifically focused on this mechanism.

Bacterial pathogens have evolved sophisticated mechanisms to manipulate host cell apoptosis, often through the various secretion systems and their effector proteins ^33^. *Stenotrophomonas maltophilia* can induce apoptosis through type II protein secretion system (T2SS) and T4SS.^34,35^ *Enteropathogenic Escherichia coli* (EPEC) triggers cell apoptosis by T3SS-secreted effectors EspF and Cif.^36,37^ In contrast, *Legionella pneumophila* T4SS effector SidF inhibits host apoptosis by directly interacting with pro-apoptotic proteins like BNIP3.^38^ Notably, observations that T6SS can induce cell apoptosis have been reported,^39,40^ but the underlying mechanisms remain unclear. The T6SS is a versatile and complex molecular machine utilized by Gram-negative bacteria to interact with their environment.^41,42^ It plays a significant role in manipulating host cells.^43–46^ T6SS also facilitates bacterial competition and our recent studies showed that these antagonistic effects can be achieved in a contact-independent manner.^47–54^ Some T6SS-secreted effectors, known as trans-kingdom effectors, function in both eukaryotic and prokaryotic cells.^55,56^ DNases associated with T6SS represent a critical class of effectors that directly cleave DNA, leading to the degradation of genetic material in target cells. This activity is highly effective in mediating interbacterial competition, as DNA degradation in competing bacteria can lead to cell death or the loss of essential genetic information, thereby providing a competitive edge to the T6SS-harboring bacterium.^51,53,54,57^ Although dozens of T6SS-secreted DNase effectors have been reported, only a limited number of studies have examined their roles in manipulating eukaryotic cellular processes. A recent study indicated that *Acinetobacter baumannii* utilizes the T6SS-secreted DNase effector TafE to target fungal cells, mediating interkingdom competition.^58^ However, the impact of T6SS DNase effectors on mammalian cells remains unknown.

In this study, we investigated the function of the T6SS in *Yersinia pseudotuberculosis*, with a focus on the trans-kingdom effector TkeA (<u>T</u>6SS-secreted trans-<u>k</u>ingdom <u>e</u>ffector inducing <u>a</u>poptosis). We demonstrated that the T6SS-secreted DNase TkeA is delivered into host cells, where it enters the nucleus and induces DNA damage. The fragmented DNA leaks into the cytoplasm, activating cGAS and leading to TNF-mediated apoptosis. Additionally, the T6SS delivers TkeA into target bacterial cells, causing genome degradation and enhancing interbacterial competition. This discovery adds a new dimension to the mechanisms employed by bacterial pathogens to manipulate host cell death pathways.

## RESULTS

### TkeA is a T6SS-3 secreted DNase effector

Gram-negative bacteria utilize the type VI secretion system (T6SS) to secrete a variety of effectors that facilitate their survival and replication in complex environments, with DNase enzymes representing a critical effector family.^51,53,54,57^ To identify new T6SS-secreted DNases, we conducted a comprehensive search of the *Yersinia pseudotuberculosis* YPIII (*Yptb*) genome, focusing on unidentified valine-glycine repeat protein G (VgrG) homologs—a structural component of T6SS often associated with adjacent effector genes. The search revealed an orphan VgrG-containing gene cluster encoding multiple potential T6SS effector-immunity pairs (*ypk_0764* to *ypk_0773*, Fig. S1A). Among these, YPK_0772 emerged as a candidate nuclease, based on its annotation as a rearrangement hotspot (Rhs) protein containing a GIY-YIG nuclease domain identified through KEGG SSDB Motif Search (https://www.kegg.jp/ssdb-bin/ssdb_motif?kid=ypy:YPK_0772).^59^

Secretion assays confirmed that YPK_0772, hereafter referred to as TkeA, is a T6SS-secreted effector (Fig. 1A). Selective inactivation of T6SS-1 to T6SS-4 by deleting ATPase ClpV1 to ClpV4 revealed that TkeA secretion is predominantly mediated by T6SS-3 (Fig. 1A). Expression of TkeA in *Escherichia coli* confirmed its toxic activity and such effect was mitigated by co-expression of the downstream immunity protein TkiA (YPK_0773) (Fig. 1B). The specific interaction between TkeA and TkiA was further validated by a bacterial two-hybrid assay (Fig. 1C). The *in vitro* DNase assay with purified TkeA confirmed its ability to degrade λ-DNA, exhibiting a degradation pattern similar to DNase I (Fig. 1D).

**Figure 1.**
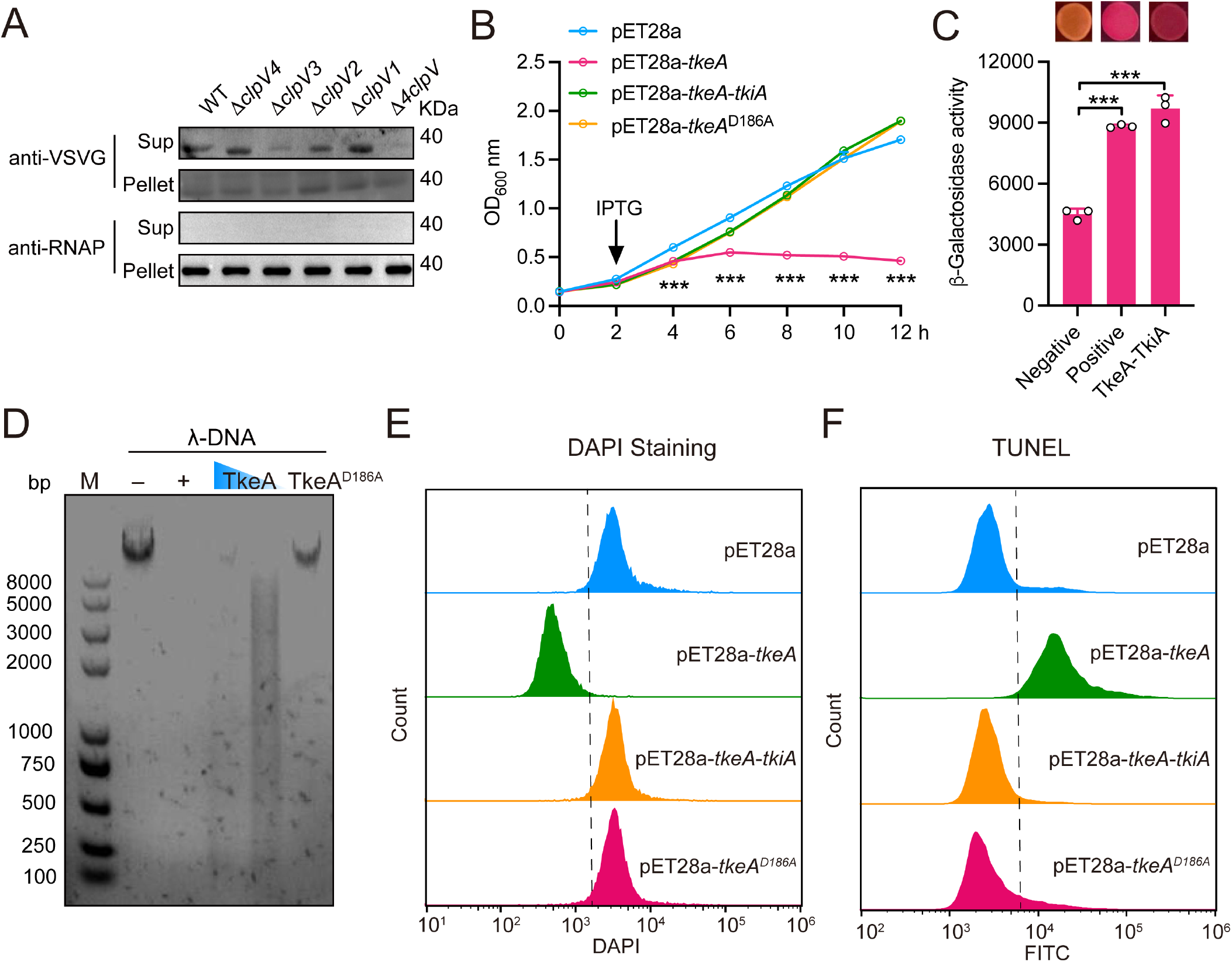
TkeA is a T6SS-3 secreted DNase effector. (A) TkeA is a T6SS3 effector. The indicated strains expressing TkeA-VSVG were cultured to OD600 = 1.6, then total cell pellets (Pellet) and secreted proteins (Sup) in the culture supernatant were isolated and probed for the presence of the TkeA protein with western blotting. Cytosolic RNA polymerase (RNAP) was used as a loading control. The Δ*4clpV* mutant is the mutant strain in which all four essential ATPase genes were deleted. The Δ*clpV1*, Δ*clpV2*, Δ*clpV3* and Δ*clpV4* is the mutant strains that deleted *clpV1*, *clpV2*, *clpV3* and *clpV4*, respectively. (B) TkeA is toxic to *E. coli*. Growth curves of *E. coli* BL21(DE3) containing indicated plasmids were determined by measuring OD600 from 0 h to 12 h at a 2 h interval. (C) Verify the interaction between TkeA and TkiA with the bacterial two-hybrid assay. Interactions were visualized with the MacConkey maltose plates (upper) and quantified with the β-galactosidase assay (lower). n = 3 (D) DNase assays indicating the integrity of λ-DNA co-incubated without (–) or with the DNase I control (+), TkeA (1 or 0.5 μM), or TkeA^D186A^ (1 μM) at 37°C for 30 min. Reaction products were analyzed using agarose gel electrophoresis. (E) Detection of the loss of DNA staining (DAPI) in indicated *E. coli* cells 4 h after IPTG induction. The *X*-axis corresponds to the 450A filter reading. (F) Detection of TkeA-induced genomic DNA fragmentation after 4 h IPTG induction in the TUNEL assay. DNA fragmentation was detected based on monitoring of fluorescence intensity (indicated on the *X*-axis) using flow cytometry. The counts resulting from cell sorting are indicated on the *Y*-axis. *P* values calculated using one-way or two-way analysis of variance (ANOVA) for multiple comparisons. Error bars represent ± SD. \*\*\**P* < 0.001. See also Figure S1.

Through random mutagenesis screening, a mutant TkeA protein (TkeA^D186A^) with loss of toxicity in *E. coli* was identified from approximately over 300 candidates (Fig. 1B). The purified TkeA^D186A^ protein lacked DNase activity, indicating that the aspartic acid residue at position 186 is critical for its enzymatic function. *In vivo* DNase activity in *E. coli* of TkeA was further corroborated by DAPI staining and Terminal deoxynucleotidyl transferase dUTP nick-end labeling (TUNEL) assays (Fig. 1E and 1F, Fig. S1B and S1C). Additionally, expression of TkeA led to around 40% of *E. coli* cells undergoing filamentation (Fig. S1B and 1D), providing additional evidence of DNA damage and halted cell division. These findings showed that TkeA is a DNase effector secreted by T6SS-3.

### TkeA causes DNA damage in mammalian cells

Since DNA is a common genetic material, it is plausible to hypothesize the effector with DNase activity is a trans-kingdom effector that targets both prokaryotic and eukaryotic cells. A recent study showed that *Acinetobacter baumannii* uses its T6SS DNase effector TafE to target fungal cells.^58^ This prompted us to investigate whether TkeA exerts toxic effects in eukaryotic cells. To verify the translocation of TkeA into HeLa cells, we fused the TEM1 (β-lactamase) reporter protein to the C terminus of TkeA and utilized *Yptb* strains expressing this fusion protein to infect HeLa cells (Fig. 2A). The T6SS-mediated transport of TkeA into HeLa cells was assessed via fluorescence resonance energy transfer (FRET) with the β-lactamase cleavable substrate CCF2/AM.^60^ Cells infected with the *Yptb* WT strain expressing TkeA-TEM1 exhibited blue fluorescence (447 nm), while cells infected with the Δ*4clpV* mutant (lacking all T6SS) predominantly displayed green fluorescence (520 nm), confirming that TkeA was effectively delivered into HeLa cells via T6SS.

**Figure 2.**
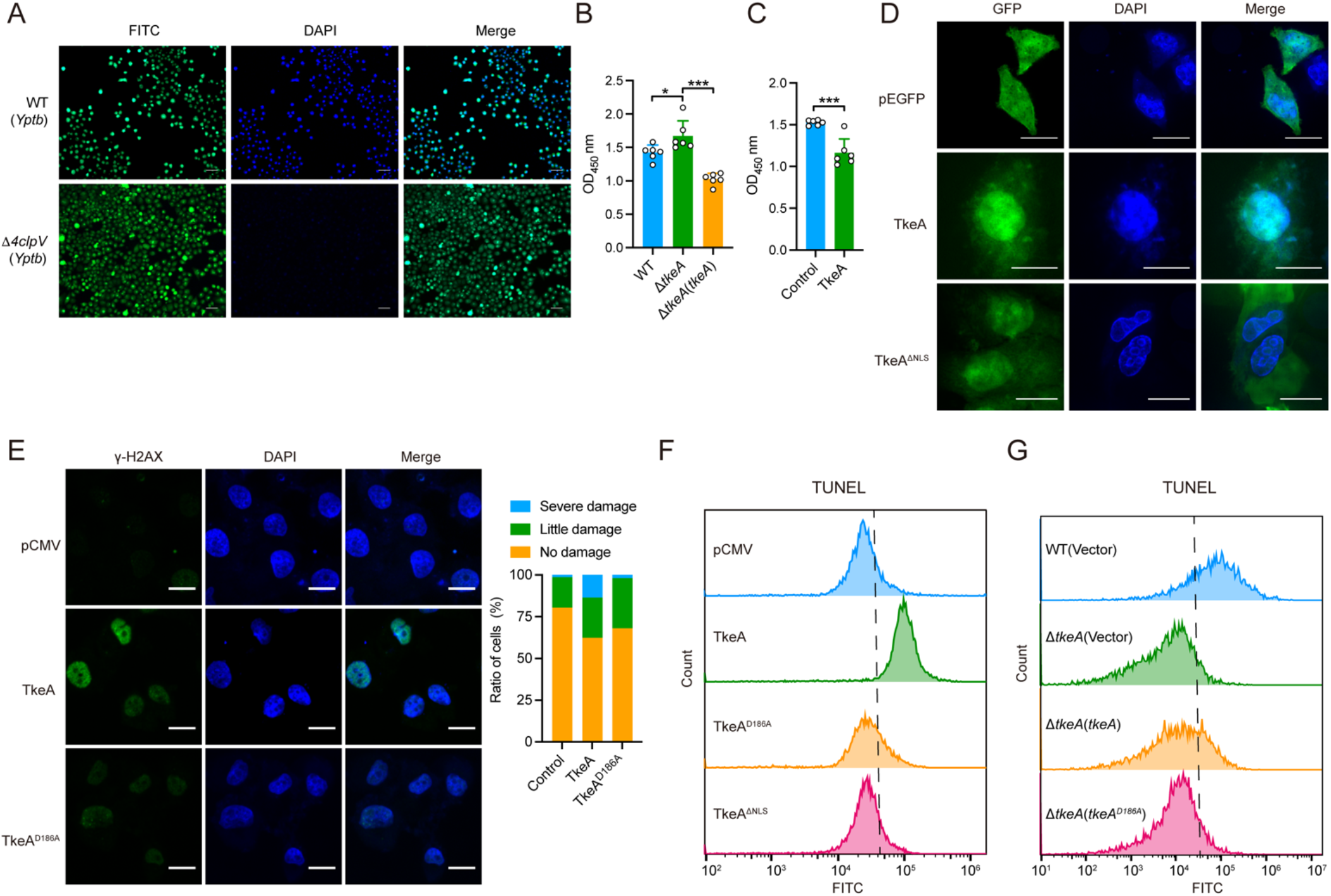
TkeA causes DNA damage in mammalian cells. (A) Translocation of TkeA in HeLa cells. HeLa cells were infected with relevant *Yptb* strains introduced with pME6032-*tkeA*-*tem* at a MOI of 100 for 1.5 h. Fluorescence microscopy was performed to visualize the translocation of TkeA. Scale bar, 50 μm. (B) The cell counting assay of the *Yptb* WT, Δ*tkeA* and Δ*tkeA* (*tkeA*) infected HeLa cells. HeLa cells were incubated with relevant *Yptb* strains at a MOI of 100 for 1.5 h. 10 μL CCK-8 was added and incubated for 4 h. Absorbance was tested at 450 nm. n = 6. (C) The cell counting assay of the TkeA-overexpressed HeLa cells. HeLa cells were transfected with pCMV (Control) or pCMV-*tkeA* vector. After 24 h, 10 μL CCK-8 was added and incubated for 4 h. Absorbance was tested at OD450 nm. n = 6. (D) Translocation of TkeA in HeLa cell. pEGFP, pEGFP-*tkeA* and pEGFP-*tkeA*^Δ*NLS*^ mutant plasmids were transfected into HeLa cells for 24 h. Florence microscopy was performed to visualize the translocation of TkeA (green). Nuclear DNA was stained with DAPI (blue). Scale bar, 200 μm. (E) Detection of DNA damage led by TkeA in HeLa cell. pCMV, pCMV-*tkeA* and pCMV-*tkeA^D186A^* were transfected into HeLa cells for 24 h. Nuclear DNA was stained with DAPI (blue). γ-H2AX signal was examined using immunofluorescence microscopy (green). The quantification was calculated on the right. Scale bar, 500 μm. (F) The indicated HeLa cells transfected with pCMV, pCMV-*tkeA*, or pCMV-*tkeA^D186A^* for 24 h by TUNEL-staining were analyzed by flow cytometry. Data are representative of three independent experiments. (G) The indicated HeLa cells infected with WT, Δ*tkeA* and Δ*tkeA* (*tkeA*) by tunnel-staining were analyzed by flow cytometry. Data are representative of three independent experiments. *P* values calculated using two-tailed Student’s *t*-test for paired comparisons or one-way analysis of variance (ANOVA) for multiple comparisons. Error bars represent ± SD. \**P* < 0.05; \*\*\**P* < 0.001. See also Figure S2.

The Cell Counting Kit-8 (CCK-8) assay was performed to evaluate the cytotoxicity of TkeA in HeLa cells. HeLa cells infected with *Yptb* Δ*tkeA* strains showed increased viability compared to those infected with the WT strain or *tkeA* complementation strain, suggesting that TkeA contributes to *Yptb*- induced cytotoxicity in HeLa cells (Fig. 2B). Additionally, HeLa cells transfected with TkeA expressing plasmid exhibited reduced viability compared to cells transfected with either an empty vector, which further confirming TkeA’s toxic effect in mammalian cells (Fig. 2C). Bioinformatic analysis identified a nuclear localization sequence (NLS: KRKKAHDRKAKK) within TkeA, suggesting that it could localize to the nucleus. This was confirmed by transfecting HeLa cells with GFP-TkeA. GFP-TkeA co-localized with nuclear DAPI staining, whereas a mutant lacking the NLS (GFP-TkeA^ΔNLS^) was primarily cytosolic (Fig. 2D). These results indicate that TkeA can localize to the nucleus of HeLa cells.

Given that CDT, a known virulence factor with DNase activity can induce DNA damage in mammalian cells,^14^ we hypothesized that TkeA might also cause DNA damage. We next assessed DNA damage by measuring the phosphorylation of γ-H2AX, a marker of DNA double-strand breaks (DSBs).^61^ HeLa cells expressing TkeA showed elevated levels of phosphorylated γ-H2AX compared to those transfected with either an empty plasmid or a plasmid harboring the TkeA^D186A^ mutant (Fig. 2E). Additionally, TkeA expression induced cell cycle arrest, further implicating its role in DNA damage (Fig. S2A and S2B). As the DNase TkeA can lead to DNA fragmentation in prokaryotic cells (Fig. 1G), we further measured the presence of DNA termini in TkeA-transfected HeLa cells. We performed a TUNEL assay in HeLa cells to assess DNA fragmentation. Cells expressing TkeA showed a significant increase in TUNEL-positive nuclei compared to controls, while cells expressing TkeA^D186A^ or TkeA^ΔNLS^ exhibited few TUNEL-positive cells (Fig. 2F). Similarly, HeLa cells infected with the *Yptb* WT strain were largely TUNEL-positive, whereas those infected with the Δ*tkeA* strain showed reduced TUNEL positivity. This reduction was reversed by complementation with *tkeA*, but not by the *tkeA^D186A^* mutant (Fig. 2G). Together, these results demonstrate that the DNase activity of TkeA induces DNA damage in mammalian cells.

### TkeA actives the cGAS-STING pathway

DNA damage often leads to DNA fragmentation that leaks into the cytoplasm.^32^ The cytoplasmic presence of these fragments is known to activate the DNA sensor cGAS, particularly following chromosomal DNA damage.^62^ To determine whether TkeA activates cGAS, we co-expressed GFP-cGAS with either TkeA or the catalytically inactive mutant TkeA^D186A^ in HeLa cells. In cells expressing TkeA, cGAS formed puncta around the nucleus, a pattern not observed in cells expressing TkeA^D186A^ (Fig. 3A). This suggests that TkeA-induced DNA damage leads to the activation of cGAS.

**Figure 3.**
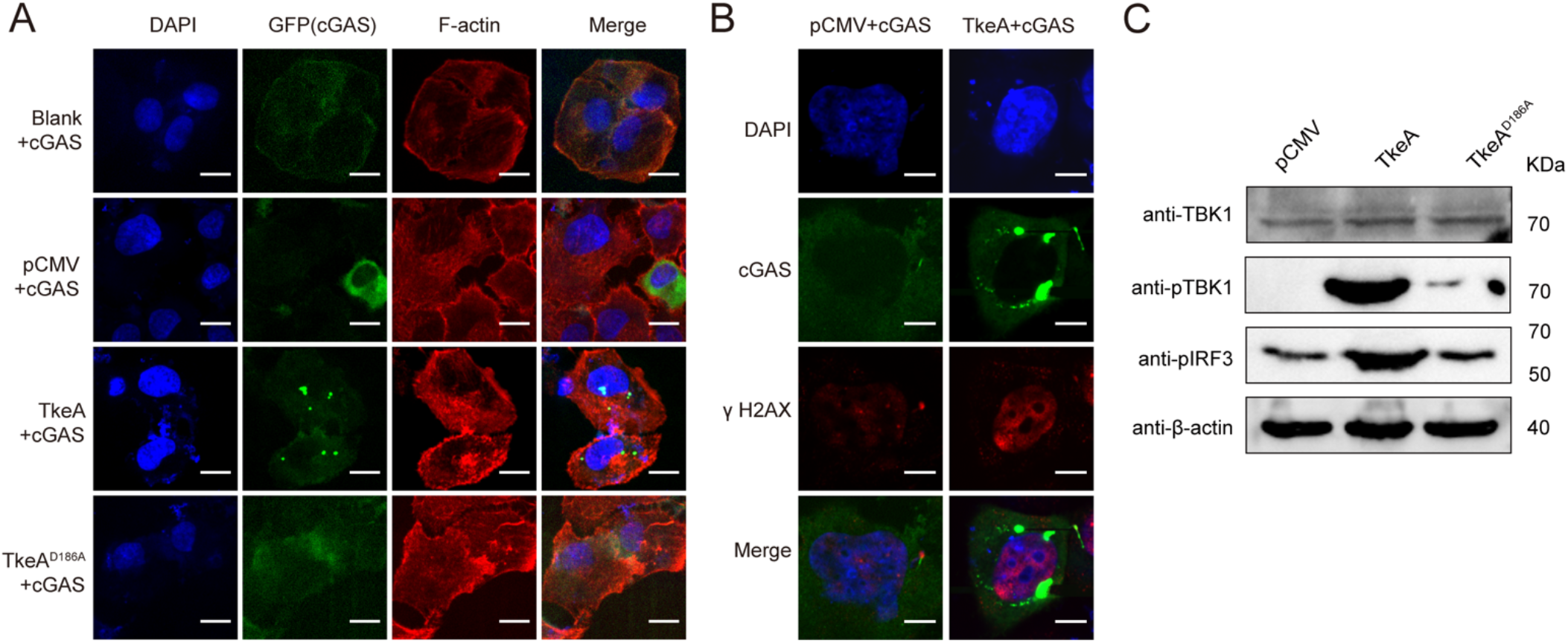
TkeA actives the cGAS-STING pathway. (A) Translocation of TkeA activates cGAS. HeLa cells were co-expressed with TkeA and cGAS-GFP for 24 h. Fluorescence microscopy was performed to visualize the activation of cGAS. DAPI, nucleus; GFP, cGAS; F-actin, cytoskeleton. Scale bar, 500 μm. (B) Representative immunofluorescence of GFP (cGAS), endogenous γ-H2AX (DNA damage) in HeLa cells to show the distribution of these proteins. HeLa cells transfected with pCMV, or pCMV-*tkeA* and cGAS-GFP for 24 h. Scale bar, 200 μm. (C) Immunoblot analysis of TBK1, phosphorylated TBK1 and IRF3 expression in HeLa cells transfected with pCMV, pCMV-*tkeA* or pCMV-*tkeA^D186A^* for 24 h.

The activation of cGAS typically involves the detection of abnormal DNA in the cytosol or its translocation from the cytoplasm to the nucleus in response to DNA damage.^32,63^ To investigate the subcellular localization of cGAS, we analyzed the distribution of GFP-cGAS, DAPI (nuclear genome), and damaged DNA (marked by γ-H2AX) in HeLa cells. The formation of cGAS-DNA foci in the cytoplasm of TkeA-expressing cells confirmed the activation of cytosolic cGAS (Fig. 3B). To further validate cGAS activation, we assessed the downstream signaling components of the cGAS-STING pathway, specifically the phosphorylation of IRF3 and TBK1 in TkeA-overexpressing HeLa cells.

Phosphorylation of both IRF3 and TBK1 was strongly induced in cells expressing TkeA, but not in those expressing TkeA^D186A^ or the control vector (Fig. 3C), confirming that TkeA activates the cGAS-STING pathway through its DNase activity.

### TkeA elicits apoptosis in a cGAS-STING-dependent manner

DNA damage is a well-known trigger for cell death and programmed cell death (PCD) including apoptosis, pyroptosis, and necroptosis.^64^ Previous studies have reported that *Yersinia* infection can induce various forms of cell death, including apoptosis and necrosis.^65,66^ To probe which PCD is elicited by TkeA, we performed Hoechst 33342/PI double staining. The presence of a brilliant red (PI) signal indicated that TkeA triggers cell apoptosis but not necrosis (Fig. 4A). Furthermore, the pan-caspase inhibitor z-VAD-FMK inhibited TkeA-induced cell death (Fig. 4B), supporting that apoptosis occurs upon TkeA transfection. The introduction of TkeA into HeLa cells also significantly increased the expression of cleaved CASPASE 3, the executioner of apoptosis (Fig. 4C). In addition, fluorescence staining further confirmed the activation of CASPASE 3 and apoptosis in TkeA-expressing HeLa cells (Fig. S3A). These findings provide compelling evidence that TkeA induces apoptosis.

**Figure 4.**
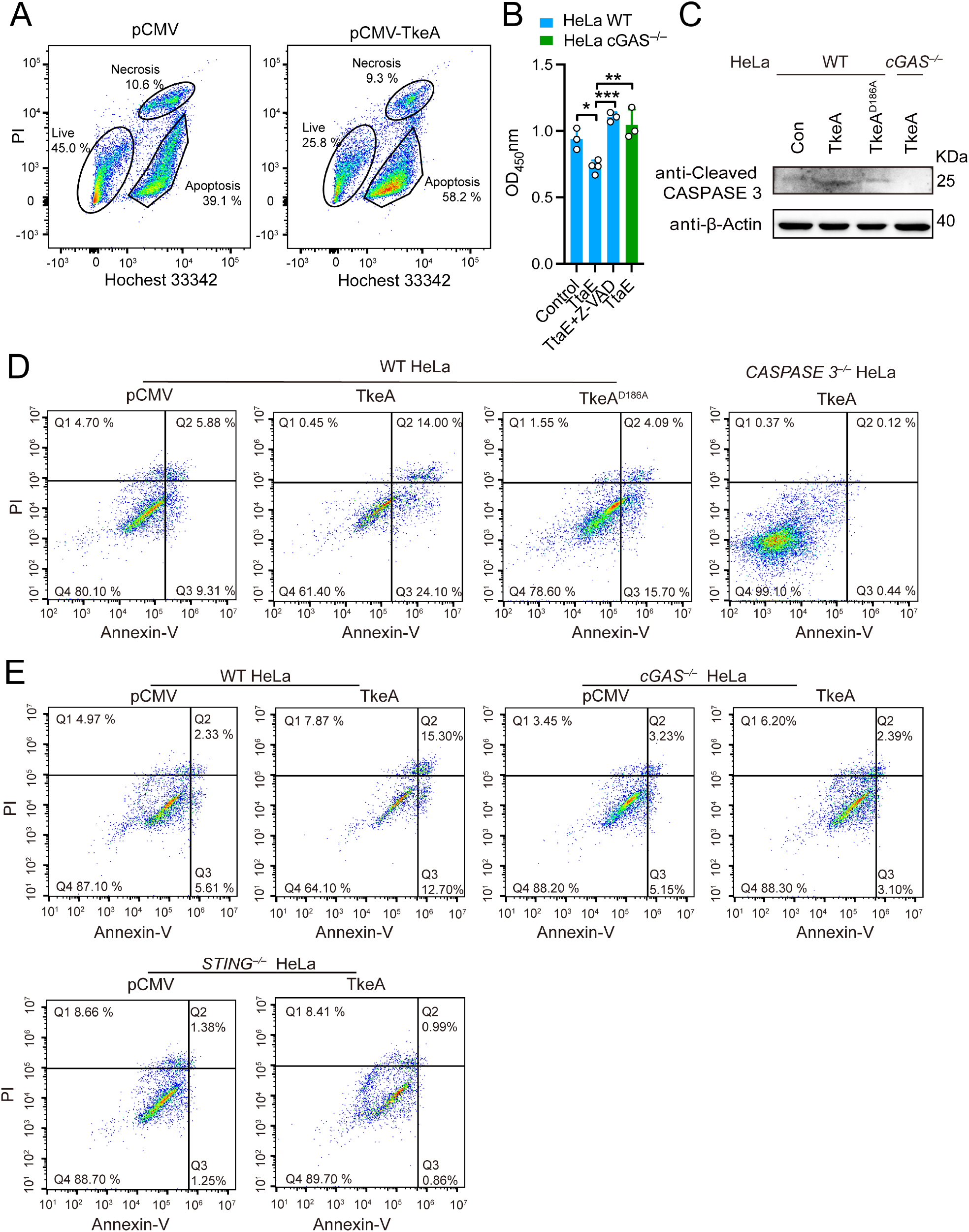
TkeA elicits apoptosis in a cGAS-STING-dependent manner. (A) HeLa cells transfected with pCMV and pCMV-*tkeA* for 24 h were collected and stained with Hoechst 33342/PI. Flow cytometry was used to identify the form of cell death. (B) The cell counting assay of the TkeA-overexpressed WT HeLa cell treated with or without z-VAD- FMK and TkeA-overexpressed *cGAS*^−/−^ HeLa cells. WT HeLa cells were transfected with pCMV (Control), pCMV-*tkeA* and pCMV-*tkeA* treated with or without z-VAD-FMK (30 μM for 6 h). *cGAS*^−/−^ HeLa cells were transfected with pCMV-*tkeA*. After 24 h, 10 μL CCK-8 was added and incubated for 4 h. Absorbance was tested at OD450 nm. n = 3. (C) Immunoblot analysis of cleaved CASPASE 3 expression in WT HeLa cells transfected with pCMV, pCMV-*tkeA* and pCMV-*tkeA^D186A^* and *cGAS*^−/−^ HeLa cells were transfected with pCMV-*tkeA* for 24 h. (D) HeLa cells transfected with pCMV, pCMV-*tkeA* and pCMV-*tkeA^D186A^*, and *CASPASE 3*^−/−^HeLa cells transfected with pCMV-*tkeA* for 24 h were collected and stained with Annexin V/PI. Flow cytometry was used to identify the cell apoptosis. (E) WT, *cGAS*^−/−^ and *STING*^−/−^ HeLa cells transfected with pCMV or pCMV-*tkeA* for 24 h and were collected and stained with Annexin V/PI. Flow cytometry was used to identify the cell apoptosis. Data in (A), (D) and (E) are from at least three biological replicates. *P* values calculated using one-way analysis of variance (ANOVA) for multiple comparisons. Error bars represent ± SD. \**P* < 0.05; \*\**P* < 0.01; \*\*\**P* < 0.001. See also Figure S3.

We further confirmed the induction of apoptosis by TkeA by Annexin V-FITC/PI double staining. Transfection of TkeA resulted in apoptosis in approximately 38.1% of cells, compared to 15.19% and 19.79% for pCMV and TkeA^D186A^, respectively (Fig. 4D). As expected, the ratio of apoptotic to non- apoptotic cells in *CASPASE 3^−/−^* HeLa cells transfected with TkeA was significantly lower than in WT HeLa cells. Furthermore, we explored the contribution of TkeA to *Yptb-*induced apoptosis by using *Yptb* WT, Δ*tkeA*, TkeA complemented strain Δ*tkeA*(*tkeA*) and catalytically inactive TkeA^D186A^ complemented strain Δ*tkeA*(*tkeA^D186A^*) to infect HeLa cells. Similarly, the levels of apoptosis in HeLa cells infected with *Yptb* WT and Δ*tkeA*(*tkeA*) were 35.0% and 27.2%, respectively, while the Δ*tkeA* and Δ*tkeA*(*tkeA^D186A^*) groups showed reduced apoptosis at 20.82% and 21.90%, respectively (Fig. S3B).

It has been reported that the leakage of DNA into the cytoplasm may activate the DNA sensor cGAS.^67^ To detect the relationship between the cGAS-STING signaling pathway and TkeA-induced cell apoptosis, we examined the cell viability in *cGAS^−/−^* HeLa cells expressing TkeA. Compared to the WT HeLa cells expressing TkeA, the cell viability of *cGAS^−/−^* HeLa cells was restored significantly (Fig. 4B), indicating that cGAS is involved in TkeA-induced apoptosis elicited by TkeA. Consistently, TkeA transfection-induced cleaved CASPASE 3 was markedly downregulated in *cGAS^−/−^* HeLa cells (Fig. 4C). Moreover, we performed the Annexin V/propidium iodide staining assay by transfecting TkeA into WT, *cGAS^−/−^* and *STING^−/−^* HeLa cells. Compared with WT HeLa cells, the TkeA-induced apoptosis was strongly reduced in *cGAS^−/−^* and *STING^−/−^* cells (Fig. 4E). Together, these results showed that TkeA elicits apoptosis in a cGAS-STING-dependent manner.

### The cGAS-STING-TNF signaling pathway is implicated in TkeA-induced apoptosis

The cGAS-STING pathway can trigger apoptosis through multiple mechanisms.^26^ To investigate the mechanisms underlying TkeA-induced apoptosis downstream of the cGAS-STING pathway, we performed RNA sequencing (RNA-seq) analysis on WT HeLa cells expressing pCMV, TkeA, and catalytically inactive TkeA^D186A^, as well as *cGAS^−/−^* HeLa cells expressing TkeA. Principal Component Analysis (PCA) showed distinct clustering for each group (Fig. S4A), and a Venn diagram highlighted the differentially expressed genes across the four groups (Fig. S4B). To identify the signaling pathways involved in cGAS-STING-dependent TkeA-induced apoptosis, we focused on genes differentially expressed in the TkeA-transfected WT HeLa group compared to other groups. In particular, those genes that downregulated in pCMV, TkeA^D186A^-transfected WT HeLa group, and TkeA-transfected *cGAS^−/−^* HeLa group, as compared to the TkeA-transfected WT HeLa group, were used to perform KEGG pathway enrichment analysis. Another criterion was gene expression that exhibited no significant difference between pCMV- and TkeA^D186A^-transfected WT HeLa cells. The most enriched pathways were the cytosolic DNA-sensing and TNF signaling pathways (Fig. 5A). Notably, many TNF signaling genes were upregulated in TkeA-transfected WT HeLa cells (Fig. 5B).

**Figure 5.**
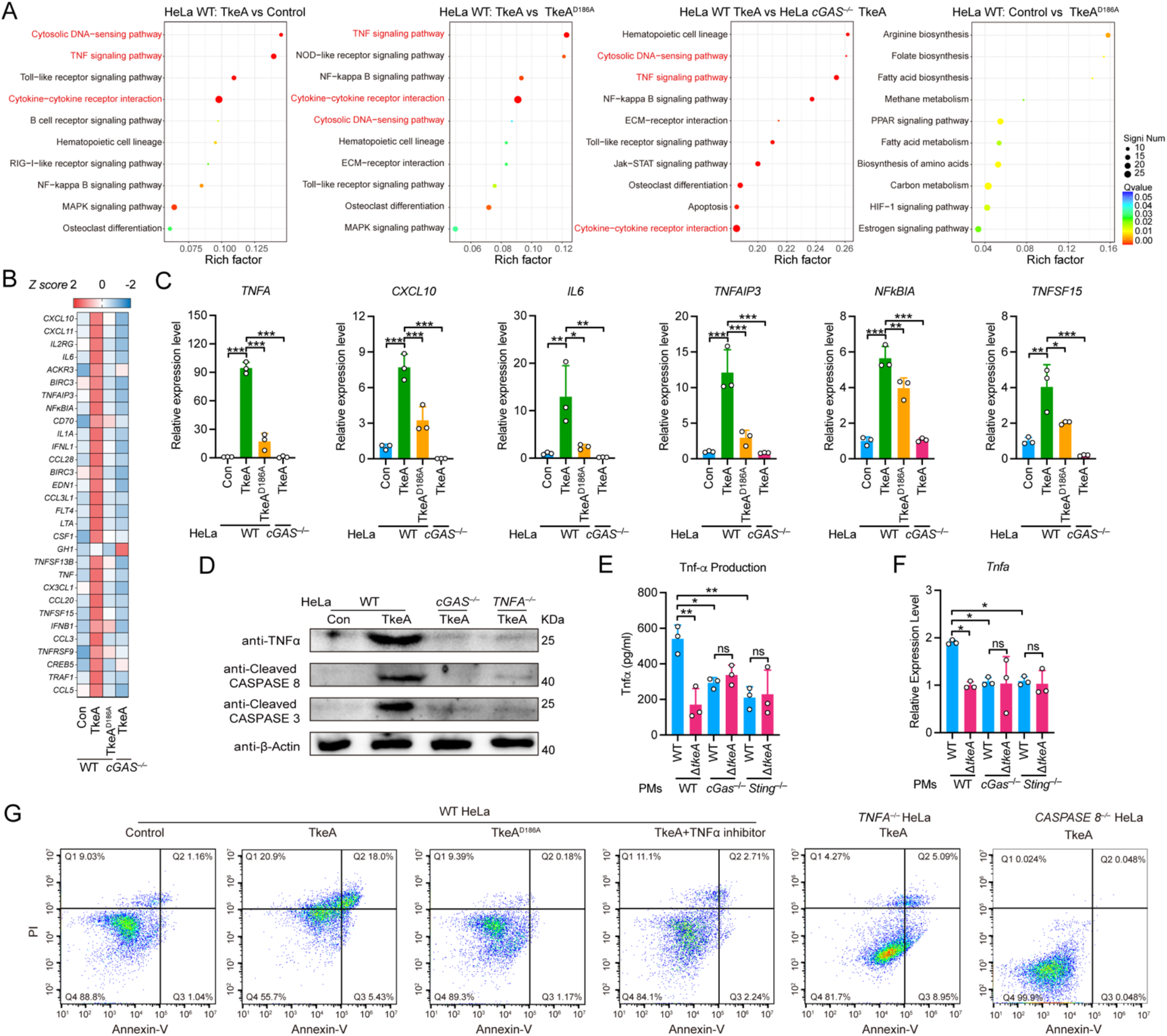
The cGAS-STING-TNF signaling pathway is implicated in TkeA-induced apoptosis. (A) Bubble chart of the top 10 significantly KEGG enriched pathways. In bubble charts, the *Y*-axis represents KEGG pathway terms; the *X*-axis indicates the Rich factor. The dot dimension corresponds to the number of genes of KEGG terms, and the dot color represents different *P* value ranges. (B) Heatmap of RNA-seq analysis which was made by calculating the RPKM. n = 3. (C) qRT-PCR analysis of gene expression in WT HeLa cells transfected with pCMV, pCMV-*tkeA* and pCMV-*tkeA^D186A,^* or cGAS^−/−^HeLa cells transfected with pCMV-*tkeA*. (D) Immunoblot analysis of protein expression in WT, *cGAS*^−/−^ and *TNFA*^−/−^ HeLa cells transfected with pCMV and pCMV-*tkeA*. (E) ELISA analysis of TNFα production in C57BL/6 wild-type, *cGas*^−/−^ and *Sting*^−/−^ mouse PMs infected with *Yptb* WT and Δ*tkeA* for 4 h at an MOI of 100. n = 3. (F) qRT-PCR analysis of gene expression in C57BL/6 wild-type, *cGas*^−/−^ and *Sting*^−/−^ mouse PMs infected with *Yptb* WT and Δ*tkeA* for 4 h at an MOI of 100. n = 3. (G) WT HeLa cells were transfected with pCMV, pCMV-*tkeA*, and 10 μM TNFα inhibitor Neochlorogenic acid was added in the pCMV-*tkeA* group. *TNFA*^−/−^ Cells or *CASPASE 8*^−/−^ cells were also transfected with pCMV-*tkeA*. All cells were collected and stained with Annexin V/PI. Flow cytometry was used to identify the cell apoptosis. *P* values calculated using one-way or two way analysis of variance (ANOVA) for multiple comparisons. Error bars represent ± SD. \**P* < 0.05; \*\**P* < 0.01; \*\*\**P* < 0.001; ns, not significant. See also Figure S4.

To validate the RNA-seq findings, we measured the expression of several TNF signaling pathway genes. TkeA transfection led to significantly higher expression of these TNF-related genes in WT HeLa cells (Fig. 5C), while the expression of *IFNB1* was not affected (Fig. S4C). To confirm that TkeA induces TNF signaling in a cGAS-STING-dependent manner, we measured TNFα expression in *cGAS^−/−^* HeLa cells following TkeA transfection. TkeA significantly increased TNFα protein levels in WT HeLa cells, but this phenotype was diminished in *cGAS^−/−^* cells (Fig. 5D). We further tested this in HeLa cells infected with *Yptb* WT and Δ*tkeA* strains. *Yptb* WT infection induced higher *TNFA* and *IL1B* mRNA expression compared to Δ*tkeA* infection (Fig. S4D). Similarly, *Yptb* WT infection induced higher TNFα secretion in mouse peritoneal macrophages (PMs), which was attenuated in *cGas*^−/−^ and *Sting*^−/−^ PMs (Fig. 5E). Of note, the expression of *Tnfa* was significantly decreased in *cGas*^−/−^ and *Sting*^−/−^ PMs (Fig. 5F). Consistently, *Yptb* Δ*tkeA* infection led to a decreased *Tnfa* expression and TNFα secretion in PMs compared to WT strain infection. Together, these results demonstrated that the cGAS-STING pathway is involved in TkeA-induced TNF production.

CASPASE 8 is a critical component of the TNF signaling pathway, responsible for activating CASPASE 3 and initiating apoptosis.^7^ To explore whether TkeA-induced activation of CASPASE 8 and CASPASE 3 is dependent on the TNF signaling pathway, we generated *TNF*^−/−^ HeLa cells. TkeA transfection significantly increased cleaved CASPASE 8 protein expression in WT HeLa cells, but this was not detected in *cGAS*^−/−^ HeLa and *TNF*^−/−^ HeLa cells (Fig. 5D). Additionally, using a TNFα inhibitor (neochlorogenic acid), we found that TkeA-induced apoptosis was markedly reduced. In addition, TkeA- induced apoptosis was diminished in *TNF*^−/−^ cells. Consistently, the deletion of *CASPASE 8* markedly suppressed TkeA-induced apoptosis (Fig. 5G). Collectively, these results demonstrate that TNF signaling is crucial for cGAS-STING-dependent TkeA-induced apoptosis.

### TkeA exerts anti-prokaryotic and anti-eukaryotic functions in mice’s gut

Successful colonization in the mouse gut requires bacteria to outcompete resident microbiota. To assess the role of TkeA in bacterial antagonism, we performed contact-dependent growth competition assays. The WT donor strain exhibited a 16-fold growth advantage over the Δ*tkeA*Δ*tkiA* recipient, which was abolished upon expressing immunity protein TkiA in the recipient strain (Fig. 6A). The antagonistic role of TkeA was further supported by interspecies competition assays. Co-incubation of *Yptb* WT with *E. coli* (Fig. 6B) or *Salmonella* Typhimurium (Fig. 6C) for 24 hours revealed the competitive advantage for *Yptb*, while deletion of *tkeA* significantly reduced this advantage.

**Figure 6.**
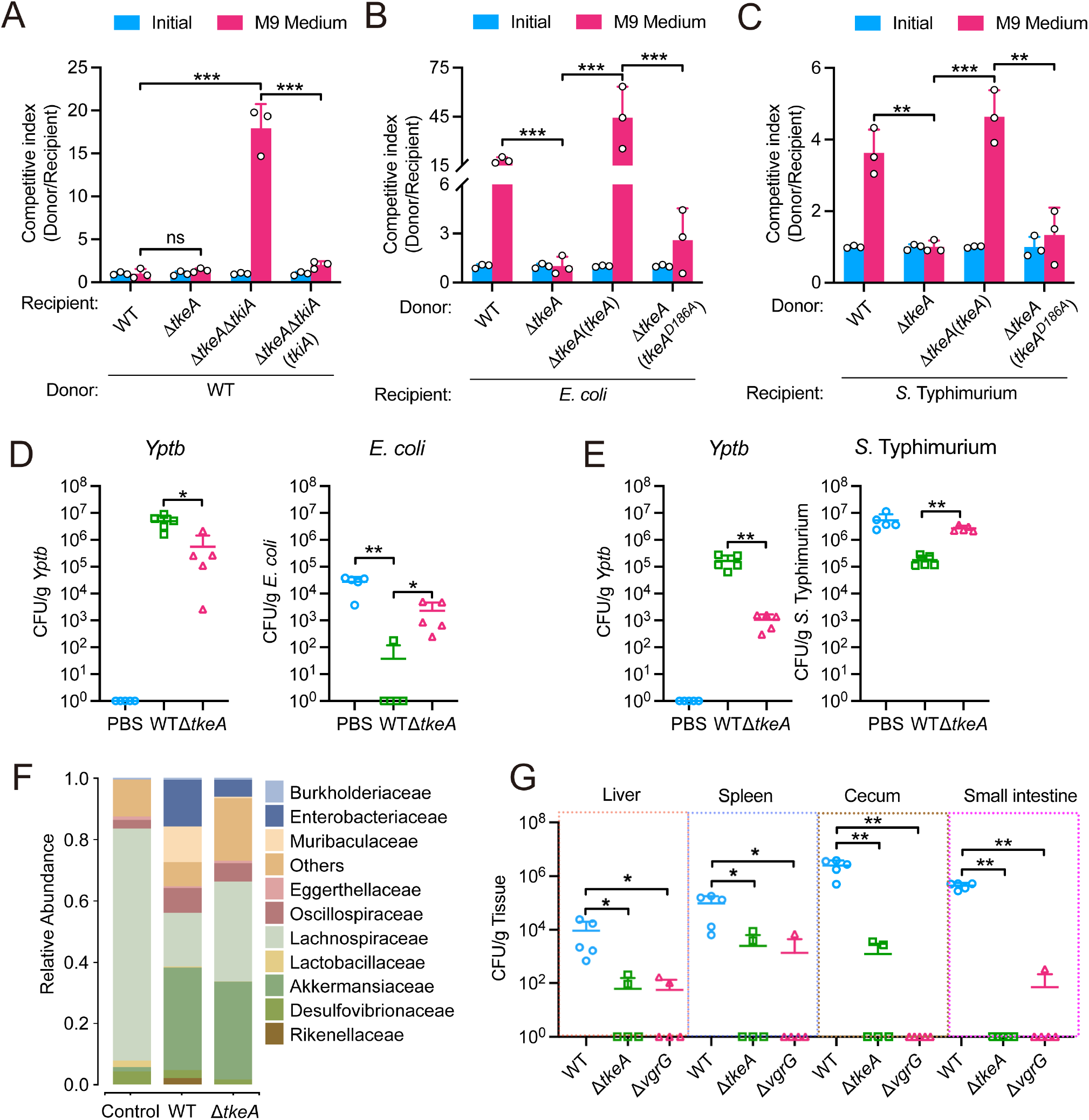
TkeA exerts anti-prokaryotic functions in mice gut. (A) Intra-species growth competition between the indicated *Yptb* donor and recipient strains. An equal amount of donor and recipient strains were mixed and grown on a solid medium for 24 h at 26 °C. The CFU ratio of donor/recipient strains was calculated based on plate counts. n = 3. (B) and (C) TkeA participates in an interference competition. Inter-species competition between the specified *Yptb* donor strains and recipient bacteria such as *Escherichia coli* (B) and *Salmonella* Typhimurium (C) in M9 liquid medium. The donor and recipient strains were combined in equal proportions and thereafter cultured for 24 h at 30°C. The *Y*-axis represents the CFU ratio of the donor and recipient strains. (D) and (E) Streptomycin-treated mice were colonized with 5×10^8^ CFUs of *E. coli* (D) or *S. Typhimurium* (E) at day 1, then challenged with 5×10^8^ CFUs of WT *Yptb*, Δ*tkeA* or PBS buffer at day 2. Animals were sacrificed on day 3, and surviving *E. coli* (D, right) or *S.* Typhimurium (E, right) and their corresponding *Yptb* strains (D, left; E, left) in the cecum were counted. PBS was used as the negative control. n = 5. (F) The analysis of the 16S rRNA gene amplicon of the cecal contents of mice that were orally gavaged with 10^9^ CFUs of different *Yptb* strains. Family-level distribution of native gut microbiota was shown under three treatments. n = 4. (G) Mice were orally gavaged with 10^9^ CFUs of different *Yptb* strains. Homogenates of the liver, spleen, cecum and small intestine were plated to determine the bacterial CFU counts per gram of organs at 24 h post-infection. n =4 - 5 *P* values calculated using the two-way analysis of variance (ANOVA) for multiple comparisons. *P* values in (D), (E), and (G) calculated using the Mann-Whitney test. Error bars represent ± SD. \**P* < 0.05; \*\**P* < 0.01; \*\*\**P* < 0.001; ns, not significant. See also Figure S5.

Complementation with WT TkeA, but not the catalytically inactive TkeA^D186A^, restored this competitive advantage, indicating TkeA’s role in *Yptb*’s fitness against *E. coli* and *S.* Typhimurium relies on its DNase activity. To test this *in vivo*, mice pre-treated with antibiotics were orally gavaged with *E. coli* (Fig. 6D) or *S.* Typhimurium (Fig. 6E) on day 1 and infected with the indicated *Yptb* strains on day 2. After 24 hours, the intestinal burden of both *E. coli* and *S.* Typhimurium was significantly reduced in mice infected with *Yptb* WT compared to those infected with Δ*tkeA*, indicating that TkeA is crucial for *in vivo* competition.

To explore TkeA’s impact on gut microbiota, we performed 16S rRNA gene sequencing on cecal contents. *Yptb* infection significantly decreased gut microbiota diversity compared to controls, with Δ*tkeA* mutant strains showing a weaker impact on taxonomic diversity than the WT (Fig. S5A). Beta diversity analysis via PCoA revealed substantial segregation in bacterial compositions among the groups (Fig. S5B). *Yptb* WT infection led to marked decreases in the phyla Verrucomicrobiota and Bacillota, with concurrent increases in Bacteroidota and Pseudomonadota (Fig. S5C). At the family level, *Yptb* WT infection resulted in decreased abundances of Eggerthellaceae, Deferribacteraceae, and Lachnospiraceae, while Enterobacteriaceae, Porphyromonadaceae, Rikenellaceae, Odoribacteraceae, Muribaculaceae, and Akkermansiaceae were increased (Fig. 6F).

Many enteric pathogens utilize the T6SS to eliminate symbionts and occupy niches within the host.^68,69^ To investigate TkeA’s role in *Yptb* colonization of mammalian organs, we orally inoculated antibiotic-pretreated and untreated mice with *Yptb* WT, Δ*tkeA*, and Δ*vgrG* strains. Colonization levels in the liver, spleen, cecum, and small intestine were measured 24 hours post-infection. The result showed that Δ*tkeA* and Δ*vgrG* strains exhibited dramatically reduced colonization compared to WT (Fig. 6G), indicating that TkeA-mediated bacterial antagonism is vital for successful colonization by outcompeting gut commensals.

### TkeA induces apoptosis in mice’s gut and contributes to the virulence of *Yptb*

To assess TkeA’s effect on *Yptb*-induced toxicity *in vivo*, we infected mice with PBS, *Yptb* WT, or Δ*tkeA* by oral gavage. Histopathological examination of cecal tissue from mice infected with Yptb WT revealed significant mucosal abscission, disorganized epithelial cell structure, and submucosal expansion (Fig. 7A), whereas mice infected with the Δ*tkeA* strain showed no discernible pathological alterations. TUNEL assays further supported these findings, with a markedly higher number of TUNEL-positive cells in the intestines of WT-infected mice compared to those in the PBS and Δ*tkeA* groups (Fig. 7B). Additionally, infection with *Yptb* WT resulted in significantly elevated mRNA levels of *Tnfa*, *Caspase 8*, and *Caspase 3* in mouse intestines, whereas these levels were substantially reduced in the Δ*tkeA*-infected mice (Fig. 7C). Correspondingly, protein analysis demonstrated that cleaved CASPASE 3 levels were higher in WT-infected mice than in those infected with the Δ*tkeA* strain, confirming that TkeA promotes apoptosis in intestinal cells (Fig. 7D). Moreover, the phosphorylation of IRF3 and TBK1—key indicators of cGAS-STING pathway activation—was observed in intestinal cells from WT-infected mice, but to a lesser extent in Δ*tkeA*-infected mice. This suggests that the cGAS-STING pathway is activated during *Yptb* infection in a TkeA-dependent manner. Finally, both Δ*tkeA*-infected and T6SS-defective Δ*vgrG*-infected mice exhibited significantly lower mortality rates compared to those infected with the WT strain (Fig. 7E). These findings demonstrate that TkeA acts as a trans-kingdom effector targeting both bacterial and mammalian cells, facilitating competition against gut bacteria and inflicting damage on intestinal epithelial cells, thus contributing to the overall virulence of *Yptb* in a mouse infection model.

**Figure 7.**
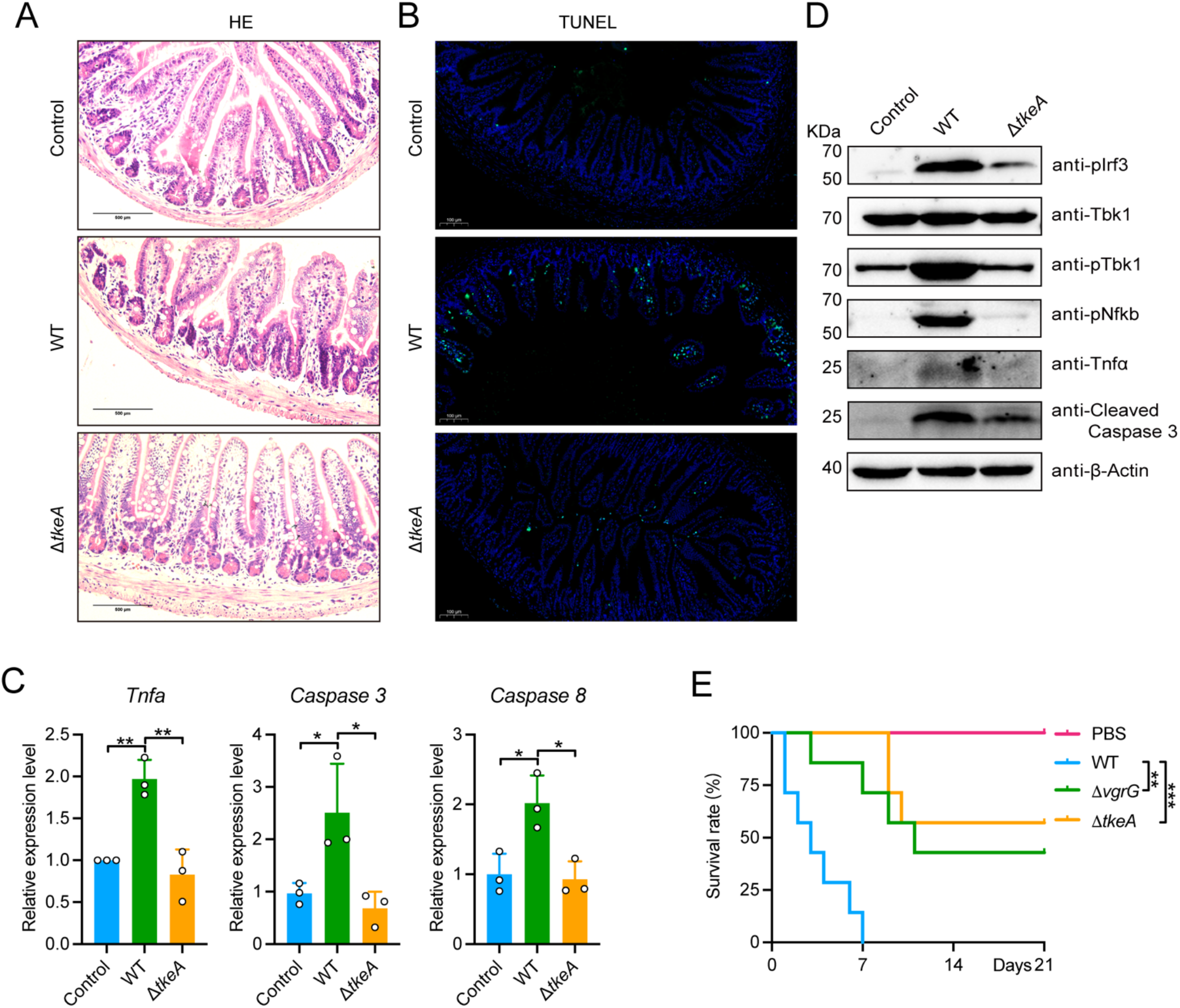
TkeA damages the intestinal epithelial cells of mice. (A) Representative microscopic pictures of H & E staining (40 ×, Scale bar, 500 μm) in mice colon infected with PBS, *Yptb* WT or Δ*tkeA* strain. (B) Representative microscopic pictures of TUNEL apoptosis assay analysis of cell apoptosis (20 ×, Scale bar, 100 μm) in mice colon infected with PBS, *Yptb* WT or Δ*tkeA* strain. (C) qRT-PCR analysis of gene expression in mice infected with PBS, *Yptb* WT or Δ*tkeA* strain. (D) Immunoblot analysis of protein expression in the small intestine from BALB/c mice infected with PBS, WT and Δ*tkeA* strain. (E) Six-week-old female BALB/c were orally gavaged with indicated *Yptb* strains which were washed by sterile PBS twice. The survival in different groups of mice was monitored daily for three weeks to establish the survival rate. n = 7. *P* values calculated using one-way analysis of variance (ANOVA) for multiple comparisons. *P* values in (E) calculated using the Log-rank test. Error bars represent ± SD. \**P* < 0.05; \*\**P* < 0.01; \*\*\**P* < 0.001. See also Figure S6.

## DISCUSSION

The T6SS has long been viewed as a potent weapon in the arsenal of Gram-negative bacteria for its ability to manipulate host cellular processes and mediate interbacterial competition. Our findings reveal a more sinister role: transforming this molecular syringe into a direct trigger of apoptosis in mammalian cells. T6SS was identified for its role in manipulating host cellular responses and the following studies revealed its role in mediating inter-bacterial competition and promoting survival in diverse ecological niches.^70^ However, our study reveals a previously unrecognized function of the T6SS, which is capable of directly inducing apoptosis in mammalian cells. This discovery is centered on a trans-kingdom DNase effector, TkeA, secreted by *Yptb*, which not only outcompete bacterial competitors by degrading their DNA but also inflicts DNA damage in host cells, triggering apoptosis through the cGAS-STING- TNF signaling pathway.

In contrast to the previously identified trans-kingdom effector TafE from *Acinetobacter baumannii*, which specifically targets fungal cells and localizes within the cytoplasm of HeLa cells,^58^ TkeA exhibits the unique capability to enter the nucleus of mammalian cells. Our observations demonstrate that GFP-labeled TkeA accumulates in the nuclei of HeLa cells, a process notably hindered when its nuclear localization signal (NLS) is deleted (Fig. 3A). This nuclear targeting by TkeA is linked to significant degradation of the host genome, support its role in DNA damage and apoptosis induction. Furthermore, while the impact of TafE on *in vivo* virulence remains unclear, our study shows that TkeA is indispensable for the virulence of *Yptb* in mouse infection models (Fig. 7E). Together, these findings showed that TkeA is a trans-kingdom effector that could disrupt the nuclei of mammalian cells to induce apoptosis.

Apoptosis is generally seen as a non-inflammatory form of programmed cell death (PCD) that cells use to prevent excessive inflammatory responses.^11^ Unlike necrosis, which is a pro-inflammatory cell death that potentially alerts the immune system,^71^ apoptosis allows bacteria to evade detection in a manner that is less likely to trigger a strong immune response. While the roles of bacterial secretion systems like T2SS, T3SS, and T4SS in inducing apoptosis have been well studied,^72–74^ the function of the T6SS in this context is not as clearly understood. Some studies suggest that T6SS can also induce apoptosis^39,40^, but the underlying mechanisms remain unclear. In this study, we showed that the T6SS effector TkeA is translocated into the nucleus of mammalian cells, where its DNase activity causes DNA damage. This damage results in the release of DNA fragments into the cytoplasm, which activates the cGAS-STING pathway. The activation of this pathway then triggers the TNF signaling cascade, leading to apoptosis through the activation of CASPASE 8. Our findings reveal a novel mechanism by which TkeA induces apoptosis, involving the cGAS-STING-TNF signaling pathway.

This mechanism is fundamentally different from how T3SS and T4SS effectors induce apoptosis. T3SS and T4SS often manipulate host signaling pathways or interfere directly with cellular machinery. For instance, T3SS effectors such as YopJ from *Yersinia* inhibit MAPK and NF-κB pathway, leading to apoptosis through the intrinsic mitochondrial pathway.^75^ Similarly, *Salmonella enterica* secretes the effector protein SipB, which interacts with Caspase-1 to cause pyroptotic cell death in macrophages.^76^ Another example is the T3SS effector IpaB from *Shigella flexneri*, which activates Caspase-1 to induce apoptosis in infected macrophages.^77^ In contrast, T6SS effectors like TkeA directly damage the host cell’s nuclear DNA, activating the cGAS-STING pathway to trigger apoptosis. In contrast, T6SS effectors such as TkeA directly target the host cell’s genetic material, inducing DNA damage that activates the DNA sensing the cGAS-STING pathway to trigger apoptosis. Notably, the apoptotic pathway triggered by TkeA is distinct from the intrinsic apoptosis pathway activated by other bacterial toxins, such as the cytolethal distending toxin (CDT). CDT primarily activates the intrinsic mitochondrial pathway through DNA damage-induced activation of p53.^14^ For example, CDT from *Aggregatibacter actinomycetemcomitans* can increase the levels of pro-apoptotic proteins Bid, Bax, and Bak in a p21CIP1/WAF1-dependent manner in Jurkat^p21−^ cells.^78^ Our findings present a new mechanism by which secretion system effectors induce apoptosis in host cells via the cGAS-STING pathway. This expands our understanding of how T6SS effectors function in the complex interactions between bacteria and host cells.

While the cGAS-STING pathway is essential for activating innate immune defenses, bacteria have evolved T6SS effectors to modulate this pathway. For instance, *Burkholderia pseudomallei* T6SS5- mediated host cell fusion triggers the cGAS-STING pathway and induces cell autophagy.^79^ In our study, we demonstrate that the T6SS effector TkeA damages nuclear DNA in host cells, leading to cGAS-STING pathway activation and subsequent apoptosis. A key aspect of TkeA in culminating apoptosis is the translocation of fragmented nuclear DNA into the cytoplasm, which then activates the cGAS pathway. Unlike the cytolethal distending toxin (CDT), which induces apoptosis via the intrinsic pathway,^14^ TkeA induces cell death through the extrinsic pathway, primarily by activating the cGAS sensor. However, the role of the cGAS-STING pathway in *Yptb* infection is complex and paradoxical.

Our previous study showed that *Yptb* employs a T6SS effector, TssS, that can chelate Mn^2+^ to counteract the STING-mediated innate immune response.^80^ This raises important questions about the contribution of the cGAS-STING pathway to *Yptb* infection progression. We speculate that the HeLa cells used in our study exhibited a minimal type I IFN response to bacterial infection, which dwarfs the *Yptb*-induced innate immune response and the induction of apoptosis becomes more prominent. The crosstalk between these pathways highlights the complexity of T6SS-induced cellular responses, where effectors can directly target host cell molecules or modulate intracellular signaling to regulate those responses.

Recent research has shown that damaged nuclear and mitochondrial DNA can induce apoptosis via the cGAS-STING pathway by activating transcription factors like IRF3 and NF-κB in HaCaT cells.^32^

Based on these findings, our study showed that TkeA-induced apoptosis is mediated through the cGAS-STING-TNF pathway. Notably, we found that the pro-apoptotic effects of TkeA are driven by TNFα activation, which leads to apoptosis. This conclusion is supported by several lines of evidence: 1) transcriptomic analysis revealed an enrichment of cytosolic DNA-sensing and TNF signaling pathways in TkeA-expressing HeLa cells; 2) higher TNFα production was observed in *Yptb* WT-infected cells compared to Δ*tkeA*-infected cells, with no difference in *cGas^−/−^* and *Sting^−/−^* PMs. Significantly lower TNFα levels were detected in the supernatant of *cGas^−/−^* and *Sting^−/−^* PMs, underscoring the critical role of the cGAS-STING pathway in mediating TNFα production; 3) inhibition of TNFα function in HeLa cells led to a reduction in apoptosis; and 4) transfection of pCMV-*tkeA* significantly affected *Tnfa* expression but had little impact on *Ifnb1* expression, indicating that TkeA induces apoptosis through TNFα rather than IFN. These results convincingly demonstrate that TNFα is the key factor mediating DNA damage-induced apoptosis. Although TNFα is primarily known as a central cytokine in inflammatory reactions, it has also been reported to induce cell death,^6^ which is not the default response of cells to TNFα. This raises the question of why TNFα preferentially induces apoptosis during *Yptb* infection. One possibility is that the TNF-TNFR1 cascades active cell death only when one of the cell death checkpoints is inactivated and those checkpoints might be inactivated during *Yptb* infection.^6,81^ However, further experiments are needed to confirm this.

Chronic activation of STING has been implicated in various inflammatory conditions, including autoimmune diseases and cancer.^25^ It is possible that DNA damage and STING activation caused by TkeA could potentially contribute to disease progression and increase the risk of carcinogenesis.

However, this hypothesis requires further investigation, particularly in the context of gut-associated bacteria and colorectal cancer (CRC). Interestingly, our study showed that WT *Yptb* strain infection in mice gut caused significant increases in the abundance of microbial families such as Enterobacteriaceae, Porphyromonadaceae, Akkermansiaceae, and Rikenellaceae—all of which are associated with an increased risk of CRC.^82–85^ Conversely, *Yptb* WT strains significantly decreased the abundances of Eggerthellaceae, Lachnospiraceae, and Lactobacillaceae, all of which are associated with a reduced risk of CRC.^86,87^ This suggests that TkeA-mediated host DNA damage, combined with alterations in gut microbiota composition, may contribute to *Yptb*-mediated intestinal carcinogenesis. However, further research is necessary to validate this hypothesis.

In conclusion, the identification of TkeA as a T6SS DNase effector that induces apoptosis through the cGAS-STING-TNF axis underscores the complexity and versatility of T6SS in bacterial pathogenesis. This study not only highlights a unique strategy employed by bacteria to manipulate host cell fate but also opens up new avenues for exploring the therapeutic potential of targeting T6SS-mediated host- pathogen interactions.

## SUPPLEMENTAL INFORMATION

Supplemental information can be found online.

## ACKNOWLEDGEMENTS

This work was supported by the grant of the National Natural Science Foundation of China (32330004 to X.S., U21A2029 to G.W., 32270134 to L.X., 31970114, 32170130 to Y.W., and 32300163 to L.S.), the Shaanxi Fundamental Science Research Project for Chemistry & Biology (Grant No. 22JHZ008 to X.S.), Ningbo Top Medical and Health Research Program (No.2023020713 to X.S.) and China Postdoctoral Science Foundation (2024T170737 to L.S. and 2024M752623 to L.S.). We thank Zhengfan Jiang (Beijing University) for generously providing the *cGas*^−/−^, *Sting*^−/−^ mice, and WT HeLa, *cGAS*^−/−^, *STING*^−/−^ cells. We also thank the Teaching and Research Core Facility at the College of Life Science (Ningjuan Fan, Xiyan Chen and Hui Duan), the Crop Biology Innovation Center at the College of Agriculture (Jianchu Zhu and Zhenzhen Ma), and Technological Innovation and Talent Cultivation Platform at College of Plant Protection (Yan Li and Zhimei Bai) for technical support.

## AUTHOR CONTRIBUTIONS

Conceptualization, L.S., L.X., X.S., and Data curation, L.S., and Formal analysis, L.S., and Funding acquisition, L.S., Y.W., L.X., G.W., X.S., and Investigation, L.S., P.Z., Z.S., Y.X., S.L., R.M., Y.W., and Methodology L.S., L.X., X.S., Project administration L.S., L.X., G.W., X.S., Resources L.S., L.X., G.W., X.S., and Supervision, G.W., X.S., and Validation, L.S., L.X., and Writing – original draft, L.S., L.X., Writing – review & editing, L.S., L.X., G.W., X.S., All authors discussed the results and commented on the manuscript.

## KEY RESOURCES TABLE

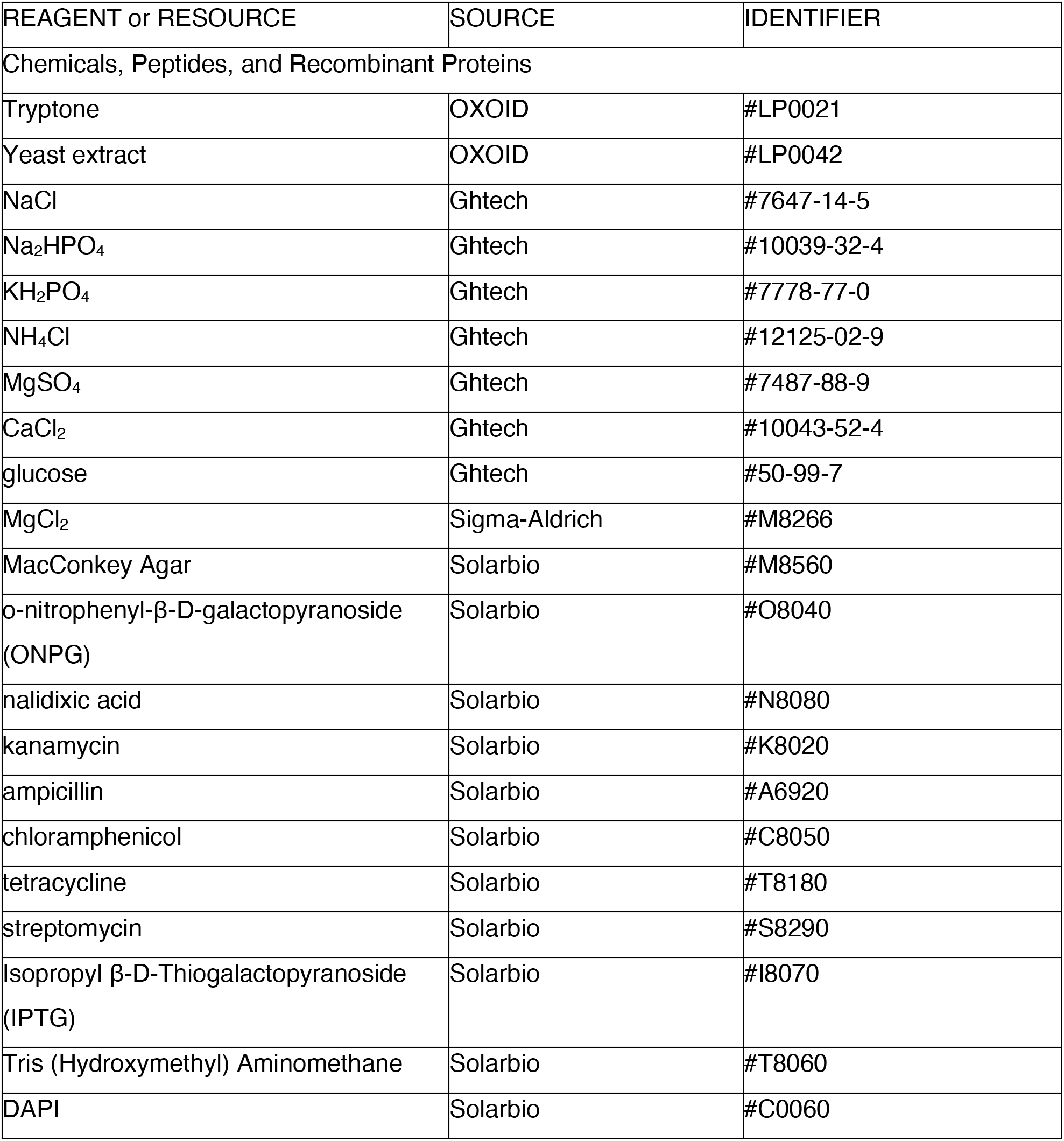

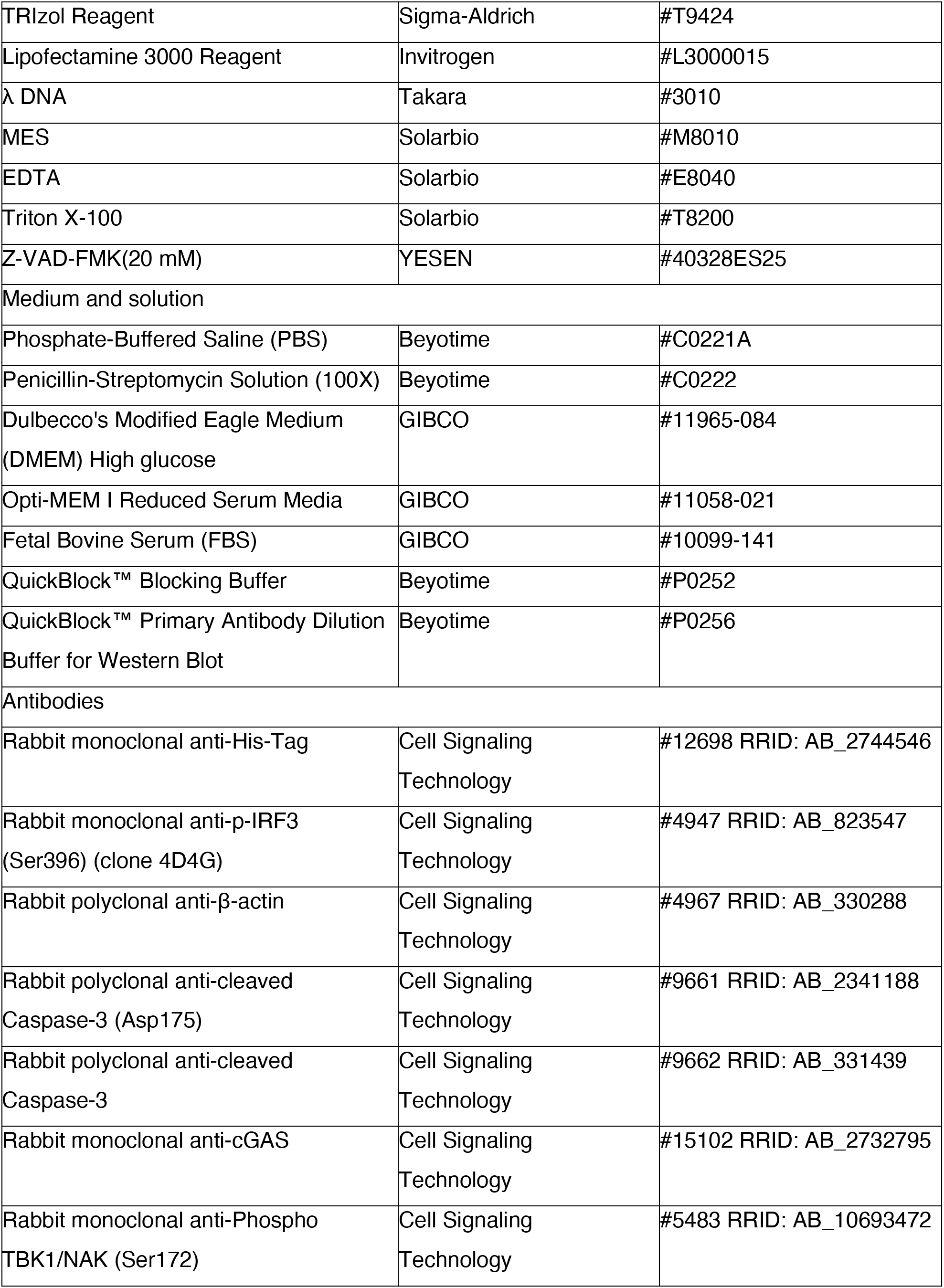

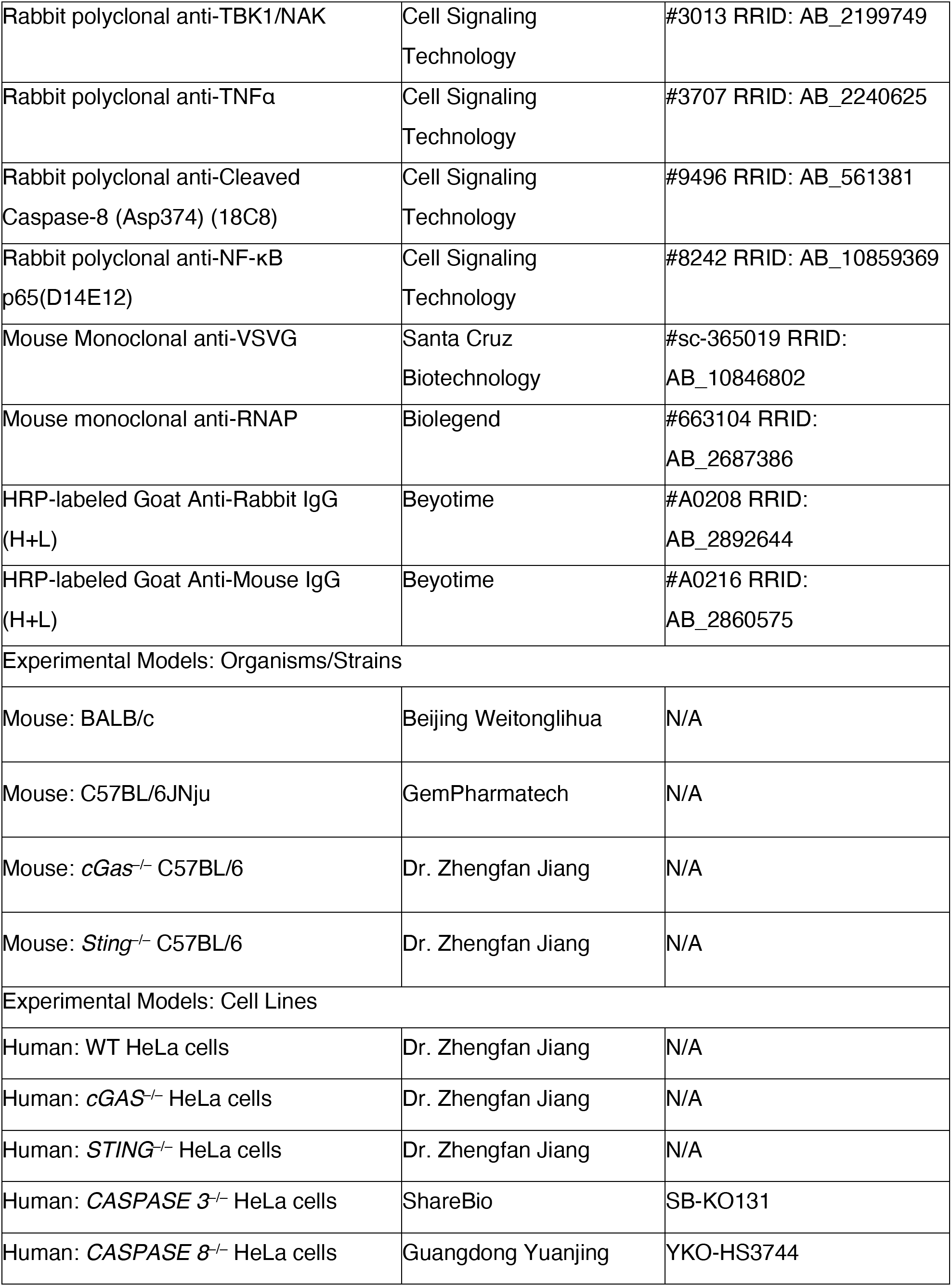

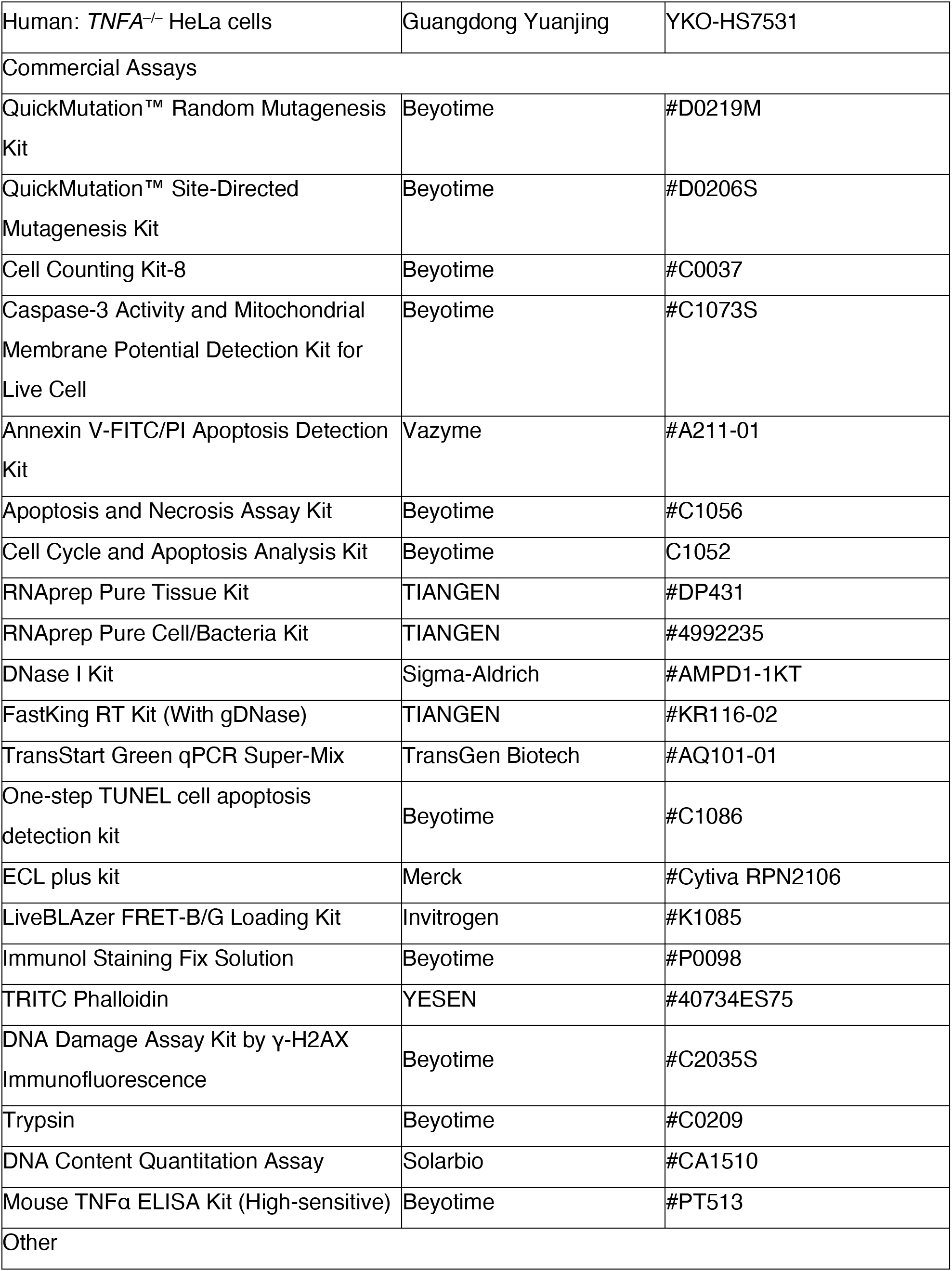

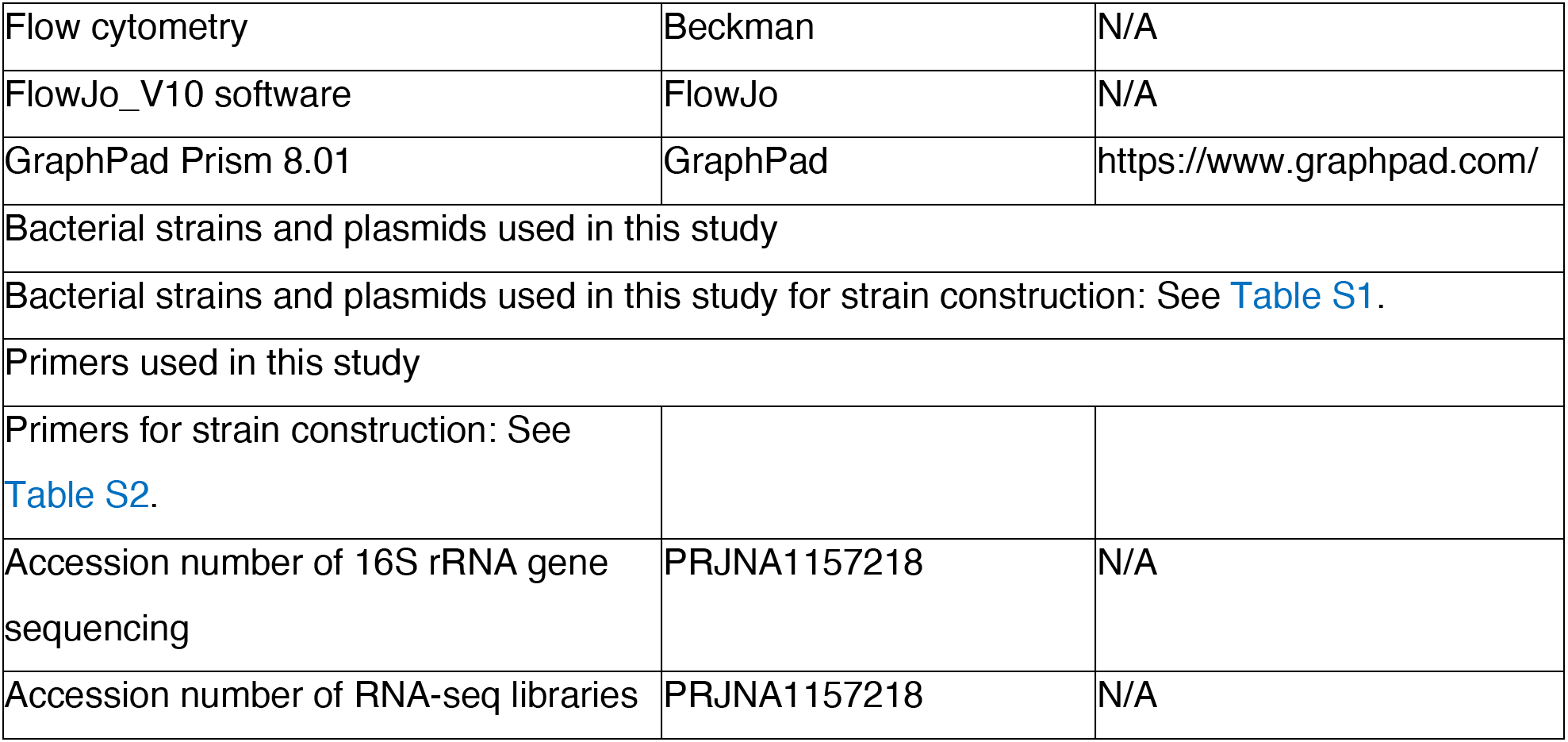

## RESOURCE AVAILABILITY

### Lead contact

Further information and requests for resources and reagents should be directed to and will be fulfilled by the lead contact, Xihui Shen.

### Materials availability

Strains and plasmids generated in this study are available upon request to the lead contact, Xihui Shen.

### Data and code availability

- ○ The 16S rRNA gene sequencing data have been deposited at the National Center for Biotechnology Information GenBank repository and China National Microbiology Data Center, and are publicly available as of the date of publication. Raw FASTQ files for the RNA-seq libraries have been deposited at the NCBI Sequence Read Archive (SRA). Accession numbers are listed in the key resources table.
- ○ This paper does not report original codes.
- ○ Any additional information required to reanalyze the data reported in this paper is available from the lead contact upon request, Xihui Shen.

## EXPERIMENTAL MODEL AND STUDY PARTICIPANT DETAILS

### Mouse studies

Six-week-old female mice (BALB/c) were purchased from Beijing Vital River Laboratory Animal Technology Co., Ltd from China. The mouse experimental procedures complied with the Regulations for the Administration of Affairs Concerning Experimental Animals, which were approved by the State Council of the People’s Republic of China. The protocol was approved by the Animal Welfare and Research Ethics Committee of Northwest A&F University (Protocol number: XN2023-1004). The mice were housed in a controlled environment with a temperature of 24 ± 2°C and a light cycle of 12 hours of light followed by 12 hours of darkness. They were provided with adlibitum access to food and water.

### Cell culture

HeLa cells were grown in DMEM media supplemented with 10% heat-inactivated FBS, 100 U/mL penicillin, and 100 g/mL streptomycin at 37°C with a CO2 concentration of 5%. Mouse peritoneal macrophages (PMs) were harvested from mice after intraperitoneal injection with beef extract peptone medium (0.3% beef extract, 1% peptone, 0.5% NaCl and 6% Soluble starch) for 3 days, and were cultured in RPMI 1640 medium, supplemented with 10% FBS, penicillin (100 U/mL), streptomycin (100 μg/mL), 10 μM sodium pyruvate, 0.1 mM non-essential amino acids, 50 mM 2-mercaptoethanol and 25 mM HEPES for 1 day.^88^

### Bacterial strains and growth conditions

Bacterial strains and plasmids utilized in this study are listed in Table S1. The *Yptb* YPⅢ strains were cultured in Yersinia-Luria-Bertani (YLB) broth (1% tryptone, 0.5% yeast extract, 0.5% NaCl) or M9 minimal medium (Na2HPO4, 6 g L^−1^; KH2PO4, 3 g L^−1^; NaCl, 0.5 g L^−1^; NH4Cl, 1 g L^−1^; MgSO4, 1 mM; CaCl2, 0.1 mM; glucose 0.2%, pH 7.0) at 26°C. *Escherichia coli* and *Salmonella* Typhimurium strains were grown in Luria Bertani (LB) with appropriate antibiotics at 37°C. The concentrations of antibiotics used in this study were as follows: nalidixic acid, 20 μg ml^-1^; kanamycin, 50 μg ml-1; ampicillin, 100 μg mL^-1^; chloramphenicol, 20 μg ml^-1^; tetracycline, 10 μg ml^-1^; streptomycin, 100 μg ml^-1^.

## METHOD DETAILS

### Plasmid construction

The primers utilized in this study are enumerated in Table S2. To acquire the expression plasmids, the genes encoding *Yptb* TkeA were amplified using polymerase chain reaction (PCR). Plasmid derivatives were created by digesting the DNA fragment and cloning it into vector pET28a with the same double restriction site. The expression clones of TkiA (YPK_0773) and TkeA-TkiA were achieved in the same manner. Overlap PCR was applied to create the plasmid pDM4-Δ*tkeA*, which was then used to create the Δ*tkeA* in-frame deletion mutant. In summary, the amplification of the upstream fragment and downstream fragment of the *tkeA* gene was carried out using the primer pairs *tkeA*-M1F-*Bam*HI/*tkeA*- M1R and *tkeA*-M2F/*tkeA*-M2R-*Sal*I. Then, both of the PCR fragments were combined using the primer pair *tkeA*-M1F-*Bam*HI/*tkeA*-M2R-*Sal*I through the process of overlap PCR. The PCR products were digested with *Bam*HI and *Sal*I, and inserted into similar digested suicide plasmid pDM4 to produce pDM4-Δ*tkeA*. The knock-out plasmid pDM4-Δ*tkeA*Δ*tkiA*, and pDM4-Δ*vgrG* were constructed with the same method.

To construct plasmids used in bacterial two-hybrid assays, the *tkeA* gene were amplified by PCR using the primer pair *tkeA*-F-*Xba*I and *tkeA*-R-*Eco*RI. Amplified DNA fragments were digested with restriction enzymes *Xba*I and *Eco*RI, and cloned into the corresponding sites of pKT25 vector. The cloning vectors pUT18C-*tkiA* were obtained with the same manner. To complement the Δ*tkeA* mutant, primers *tkeA*-F- *Spe*I and *tkeA*-R-*Sal*I were employed to amplify the *tkeA* gene fragment from *Yptb* genomic DNA. The PCR product was digested with *Spe*I/*Sal*I and ligated into similarly digested pKT100 to produce pKT100-*tkeA*. The complementary plasmids pKT100-*tkiA*, pKT100-*vgrG*, pEGFP-*tkeA* and pCMV-*tkeA* were constructed similarly. Plasmid pME6032-*tkeA*-vsvg was constructed for protein secretion assay. Briefly, primers *tkeA*-F-*Eco*RI and *tkeA*-R-*vsvg*-*Bgl*II were employed to amplify the *tkeA* gene from *Yptb* genomic DNA. The PCR product was digested with *Eco*RI/*Bgl*II and inserted into similarly digested pME6032 to generate pME6032-*tkeA*-*vsvg*. To construct TEM1 translocation reporter vector, primers *tkeA*-F-*Eco*RI/*tkeA*-R-*Bgl*II were utilized to amplify the gene *tkeA* from the *Yptb* genome DNA. Then, the fragment *tkeA* was inserted into the pME6032-*tem1* with the same digested sites to produce pME6032-*tkeA*-*tem1*. The primer pairs *tkeA^D186A^*-F and *tkeA^D186A^*-R were used to amplify the complete plasmid pET28a-*tkeA^D186A^*, pKT100-*tkeA^D186A^*, pEGFP-*tkeA^D186A^* and pCMV-*tkeA^D186A^* using QuickMutation™ Site-Directed Mutagenesis Kit. To obtain pCMV-*cGAS*-*GFP*, primers *cgas*-F- *Hin*dIII/*cgas*-R-*Bgl*II were used to amplify the fragment *cGAS* of human. Then, *cGAS* were inserted into pCMV-C-EGFP with the same digested sites *Hin*dIII/*Bgl*II. The integrity of the insert in all constructs was confirmed by DNA sequencing.

### Overexpression and purification of recombinant proteins

In order to achieve the expression and purification of recombinant proteins tagged with His6 and GST, the plasmid pET28a derivatives were introduced into *Escherichia coli* strains BL21(DE3). The bacteria were cultured in 5 mL of Luria-Bertani (LB) medium at 37°C until reached the stationary phase. Then bacteria were re-inoculated into fresh LB medium at a ratio of 1:100 and cultivated at 37°C until the optical density at 600 nm (OD600) reached a value of 0.40. Subsequently, a concentration of 0.2-0.5 mM IPTG was added into the growth medium, and the cultivation process was extended for an additional 12 hours at 16°C with150 rpm. The cells were harvested and subjected to sonication for disruption.

Purification was carried out using either the His•Bind Ni-NTA resin (Novagen, Madison, WI), following the instructions provided by the manufacturer. The purified proteins underwent dialysis against phosphate-buffered saline (PBS) at 4°C for overnight.

### Protein secretion assay

To conduct the secretion assay, *Yptb* strains were cultivated in 3 mL of YLB medium at 26°C. Subsequently, the cultures were transfected into 300 mL of YLB medium supplemented with 1 mM IPTG until OD600 reached 1.60. A total volume of 2 mL of culture solution was obtained, and cell pellets were resuspended in the SDS-PAGE sample loading buffer. A volume of 280 mL of culture medium was centrifuged at 5,000 rpm for 20 minutes. Next, the supernatant was centrifuged at a speed of 9,900 rpm for another 50 minutes. The final supernatant was collected and filtered with a 0.22 μm pore size filter (Millipore, MA). The proteins were collected by filtration with a nitrocellulose filter three times (BA85, Whatman, Germany). The filter was dissolved in 100 μl of SDS loading buffer and incubated at 65°C for 15 minutes, then boiled for 10 min to recover the protein present. The protein samples from the total cell pellet and culture supernatant were separated using SDS-PAGE and subsequently analyzed through the Western blot.

### Bacterial two-hybrid assay

The bacterial two-hybrid complementation assays were conducted following the methods outlined in previous studies. In this study, the pKT25 and pUT18C derivatives were co-transformed into *E. coli* BTH101. The transformed cells were then cultured on a MacConkey plate supplemented with Ampicillin (100 μg ml^-1^), Kanamycin (50 μg ml^-1^), and IPTG (1 mM) at 30°C. Simultaneously, the plasmid pKT25- zip/pUT18C-zip and pKT25/pUT18C were introduced into *E. coli* BTH101, with the former serving as the positive control and the latter as the negative control. The interactions were assessed by employing the MacConkey medium, whereby the presence of a red colony color indicates a protein interaction.

The quantification of protein interactions was conducted by measuring the activities of β-galactosidase in liquid cultures. In summary, overnight cultures were diluted to a concentration of 1% and subsequently cultivated in LB broth supplemented with antibiotics at 30°C until the OD600 reached 1.0.

The enzymatic activity of β-galactosidase was then evaluated using o-nitrophenyl-β-D- galactopyranoside (ONPG) as the substrate.

### Growth inhibition assay

*E. coli* BL21(DE3) strains containing pET28a, pET28a-*tkeA*, pET28a-*tkeA^D186A^*, and pET28a-*tkeA*-*tkiA* plasmids were cultured in an LB medium. The cultures that had been incubated overnight were standardized to achieve the same optical density and subsequently diluted by a factor of 100 into LB broth supplemented with suitable antibiotics. Following incubation at 26°C and 180 rpm for 2 hours, the expression of recombinant proteins was added with 0.5 mM IPTG. Subsequently, the incubation was sustained under the same condition. The monitoring of cultural growth was conducted by measuring OD600 at regular intervals of 2 hours.

### Western blot analysis

The protein samples underwent resolution through SDS-PAGE and subsequent transfer onto PVDF membranes (Millipore, MA). Subsequently, the membrane was blocked using a 5% (w/v) BSA solution for 8 hours at 4°C. Following this, the membrane was incubated with primary antibodies overnight at 4°C. The antibodies used in this study were as follows: anti-VSVG at a dilution of 1:1000; anti-RNAP at a dilution of 1:400; and anti-His at a dilution of 1:500. The rest of the antibodies were used at a dilution of 1:1000. The membrane was washed in TBST buffer, consisting of 50 mM Tris, 150 mM NaCl, 0.05% Tween 20, and pH 7.4. Subsequently, it was incubated with secondary antibodies for 4 hours at 4°C. Following the incubation, the membrane was washed for 5 times using TBST buffer. The signals were detected by employing the ECL plus kit in conjunction with a Chemiluminescence imager (Tanon 5200Multi, Beijing).

### Construction of mutant library by epPCR

The plasmid pET28a-*tkeA* was subjected to error-prone PCR (epPCR) using the QuickMutation™ Random Mutagenesis Kit. The primers *tkeA*-F-*Eco*RI and *tkeA*-R-*Sal*I were utilized according to the manufacturer’s instructions. The experimental polymerase chain reaction (epPCR) program was implemented in the following process: 94 °C for 3 min, 30 cycles of 30 s at 94 °C, 30 s at 55 °C, and 30 s at 72 °C, followed by 10 min at 72 °C final extension. The amplified DNA fragments obtained from PCR were subjected to gel purification. Subsequently, these purified fragments were digested using *Eco*RI and *Sal*I. The resulting digested fragments were then inserted into pET28a plasmids that had been similarly treated with *Eco*RI and *Sal*I enzymes. The ligation mixture was transformed into BL21(DE3).

Transformants lost toxicity were screened in an LB medium containing 0.3 mM IPTG and were further verified by cloning the mutated alleles of *tkeA* into a new vector. The mutations were identified through the process of DNA sequencing analysis.

### DNase assay

The TkeA protein purified was incubated with λ DNA in a reaction buffer consisting of 20 mM MES, 100 mM NaCl, 2 mM MgCl2, and pH 6.9. In all, 4 mM EDTA was supplemented in the reaction system. The DNA hydrolysis was conducted at 37°C for 30 minutes. The state of DNA integrity was subsequently assessed using 0.7% agarose gel electrophoresis.^53,54^

### DAPI staining and flow cytometry analysis for bacteria

The DAPI staining and flow cytometry analysis were conducted according to the methods described before. The *E. coli* BL21(DE3) strain, harboring the pET28a plasmid or its derivatives expressing TkeA alone (pET28a-*tkeA*) or TkeA-TkiA together (pET28a-*tkeA*-*tkiA*), was cultured overnight. Subsequently, the culture was diluted with the ratio of 1/100 into LB broth and incubated at 37°C with 180 rpm. After 2 hours, the samples were stimulated through the introduction of 0.5 mM IPTG. Subsequently, cultivation was sustained for an additional 4 hours at 26°C. The collected cells were washed by phosphate- buffered saline (PBS). The cells were fixed and incubated in a solution containing 0.3% Triton X-100 in PBS for 5 minutes. Following this, the cells were stained using a concentration of 10 μg ml^-1^ of DAPI for 5 minutes at 37°C. The stained cells were then washed with PBS three times. Finally, the cells were examined using either a fluorescence microscope (Andor Revolution-XD, Britain) or flow cytometry (Beckman, CytoFLEX). A total of 10,000 cells were collected for each sample and analyzed using FlowJo_V10 software.

### Quantitative Real-time PCR (qRT-PCR)

Exponentially growing strains were subjected to total RNA isolation using the RNAprep Pure Cell/Bacteria Kit in conjunction with the DNase I Kit. The concentration of RNA was determined by the NanoDrop 2000 spectrophotometer (Thermo Fisher Scientific, USA). The measurement of mRNA abundance in each of the samples was conducted by the TransStart Green qPCR Super-Mix and the Bio-Rad CFX96 Real-Time PCR Detection System (Bio-Rad, USA), following the instructions provided by the manufacturers. The primers used in this investigation are listed in Table S2. To standardize the results, the internal standard of relative abundance of 16S rRNA was employed.

### Intra-species and inter-species competition *in vitro*

The intra-species competition assays were conducted following the method described with slight modification. In summary, strains grown overnight were washed and adjusted to an optical density of 1.0 at OD600 using an M9 medium. These adjusted strains were then combined to conduct a competition. The donor-to-recipient ratio at the beginning of the experiment was 1:1. The co-cultures were then introduced into LB/M9 medium and incubated at 30°C for 24 hours. To perform inter-species competition assays, *Yptb* strains and *Escherichia coli* (DH5α) or *Salmonella* Typhimurium strains were cultured overnight. The resulting cultures were washed three times using M9 liquid and subsequently adjusted to an optical density of 1.0 at OD600. The *Yptb* strains and target strains were combined in equal proportions and subjected to incubation at 30°C for 24 hours. The CFU ratio of the donor and recipient strains was assessed through plate counts at corresponding time intervals subsequent to the competition. The data obtained from all competitions were analyzed using the one-way or two-way ANOVA test. The presented results represent the average of a single representative assay that was conducted three times.

### Mouse infection

Mid-exponential phase *Yptb* strains were grown in YLB medium at 26°C, washed with saline water twice, and then concentrated to 12.5-bold. Six-week-old female mice were orally gavaged with 100 μL indicated *Yptb* strains with 10^9^ CFU, and the survival rate of the mice was measured daily for 21 days.

### Murine colonization assay

Six-week-old female BALB/c mice were adapted in the laboratory for three days. Subsequently, they were orally gavaged with 10^9^ CFUs of the corresponding *Yptb* strains. The mice were then observed and monitored for either 24 or 48 hours. At the end of the experiment, mice were sacrificed, and the liver, spleen, cecum, and small intestine tissue were ground and plated on selective YLB antibiotic plates for CFU enumeration.

### Translocation assay for TkeA::TEM1 fusions

The translocation experiment was carried out as previously described. TEM1 fusion TkeA expressing bacterial strains were incubated with HeLa cells (at a MOI of 100) for 1.5 h in 96-well black-wall, clear- bottom plates. After three washes in PBS, HeLa cells were treated for 90 min at room temperature with CCF2-AM (LiveBLAzer FRET-B/G Loading Kit). Fluorescence was measured using a microtiter plate reader at an excitation wavelength of 410 nm following the manufacturer’s instructions. Translocation was shown using a comparison of the cleaved (blue, 450 nm) and uncleaved (green, 520 nm) signals. A Nikon fluorescent microscope (Nikon, Japan) was used to look at the materials up close and personal.

### CCK-8 assay

Cell toxicity experiments were conducted following the guidelines provided by the manufacturer of Cell Counting Kit 8 (CCK8). HeLa cells were inoculated into 96-well plates and then exposed to the specified *Yptb* strains (MOI=100) or transfected with pCMV, TkeA, or TkeA^D186A^ constructs for 24 hours. The cells were stained using a regnant solution in the kit with a concentration of 10% (v/v). After 30 min, the absorbance at 450 nm was quantified using a microplate reader.

### Fluorescence assay and Immunofluorescence assay

Cells were seeded on Glass Bottom Cell Culture Dish (Biosharp) and were transfected with the indicated plasmid for 24 h. Treated cells were washed with PBS buffer twice and fixed with Immunol Staining Fix Solution for 15 min at room temperature. The fixative was removed, and the cells were washed three times with PBS Buffer for 3-5 min each wash. As for the fluorescence assay, the cells were incubated with PBS containing 0.3% Triton X-100 for 5 min, and stained using TRITC Phalloidin. The nuclear stain (DAPI) was added for 5 min and cells were washed 3 times. As for the Immunofluorescence assay, the DNA Damage Assay Kit by γ-H2AX Immunofluorescence was used to do the following steps. Briefly, after blocking the cells for an hour, γ-H2AX Rabbit mAb was added to incubate for 1 h at room temperature. Cells were washed 3 times with PBS for 5-10 minutes each time. Next, after incubating with anti-rabbit 488 or anti-rabbit 555 for 1 h, the cells were washed twice with PBS for 5-10 minutes each time. DAPI was added for 5 min and cells were washed 3 times. γ-H2AX staining exhibits green or red fluorescence and DAPI staining of nuclei exhibits blue fluorescence.

Images were acquired via a high-speed rotary disc-type fluorescence confocal microscope (Andor Revolution-XD, UK).

### TUNEL assay for HeLa cells

HeLa cells were seeded into a 24-well plate and transfected with an indicated plasmid for 24 h. Then, cells were collected by 0.25% Trypsin and washed with PBS. Next, the collected cells were fixed, incubated with PBS containing 0.3% Triton X-100 for 5 min, and stained using the One Step TUNEL Apoptosis Assay Kit. Flow cytometry (Beckman, CytoFLEX) was used to detect the fluorescence intensity. 10000 cells were gathered for each sample and analyzed by FlowJo_V10.

### CASPASE 3 activity and apoptosis detection for live cells

The CASPASE 3 Activity and Apoptosis Detection Kit for Live Cell was used to do this experiment. In short, the treated cells transfected with the indicated plasmid were incubated with a staining solution containing Annexin V-mCherry and 1mM GreenNuc™ CASPASE 3 Substrate for 30 min. Images were acquired via a high-speed rotary disc-type fluorescence confocal microscope (Andor Revolution-XD, UK).

### RNA-Seq experiment

Whole transcriptome sequencing was conducted at Sangon Biotech (Shanghai, China). Following TRIzol-mediated total RNA isolation from the cells, we examined the RNA for signs of degradation and contamination on 1% agarose gels, determined the RNA’s purity using a NanoPhotometer spectrophotometer (Implen), and evaluated its integrity with a Bio-analyzer 2100 system. The sequencing procedures were carried out in the same way as previously reported. Magnetic beads with Oligo(dT) attached were utilized to purify mRNA. The procedures of cDNA synthesis, end repair, A- base addition, and ligation of the Illumina-indexed adaptors were carried out in accordance with the provided instructions. The final library was made by denaturing and circularizing the double-stranded PCR products from the previous step with the splint oligo sequence. The final library was prepared by formatting the single-strand circular DNA (ssCir DNA). In order to create the DNA nanoball (DNB), the final library was amplified using phi29. In this study, we used the BGIseq500 platform to obtain single- end 50-base reads from DNBs placed into a patterned nanoarray. DESeq2 was used for the differential expression analysis and a threshold was set to a Q value with a false discovery rate (FDR) < 0.05.

### Transfection

The Lipofectamine 3000 Reagent was utilized for the purpose of transfecting DNA into HeLa cells. The cells were seeded into 12-well plates or 24-well plates with a density of 2 × 10^5^ cells per well. On the following day, the cells were subjected to transfection using plasmid DNA.

### HeLa cell infection

Grown-overnight *Yptb* strains were cultured in YLB at 30°C with appropriate antibiotics. The next day, the pellets were collected and resuspended in PBS. HeLa cells were grown in DMEM devoid of FBS and penicillin-streptomycin prior to infection. HeLa cells were infected with *Yptb* strains at a MOI of 100. After centrifuging at 500 g for 5 minutes to bring the bacteria closer to the cells, the infected cells were incubated at 37°C for 2 hours. Then the cells were washed with PBS twice and resuspended in FBS- and penicillin- and streptomycin-containing media and cultured at 37°C in 5% CO2 for 2 h. At a total of 4 hours’ infection, cells were taken out for RNA isolation and cell toxicity test.

### Hochest 33342/propidium iodide (PI) assay

Apoptosis and Necrosis Assay Kit was used to detect the form of cell death according to the manufacturer’s protocol. In short, 5 × 10^5^ HeLa cells transfected with the specified plasmid were collected and washed twice with PBS solution. Cells were suspended in 100 μl of Binding Buffer and then treated with Hochest 33342 and PI Staining Solution for 30 minutes on ice. Flow cytometry (Beckman, CytoFLEX) was used to examine the cell’s condition.

### Annexin V-FITC/propidium iodide (PI) assay

Annexin V-FITC/PI Apoptosis Detection Kit was used to detect cell apoptosis following the manufacturer’s protocol. Briefly, a total of 5 × 10^5^ HeLa cells transfected with the indicated plasmid were collected and washed twice with PBS buffer. Cells were resuspended in 100 μL 1 × Binding Buffer and incubated with Annexin V-FITC and PI Staining Solution for 10 min in the dark. Flow cytometry (Beckman, CytoFLEX) was utilized to analyze the apoptotic cells.

### Cell cycle analysis

The distribution of treated HeLa cells in the cell cycle phases was determined by measuring DNA content using DNA Content Quantitation Assay. In brief, treated cells were collected by 0.25% trypsin and fixed with the ice-cold 70% ethanol overnight at 4 °C. Subsequently, cells were centrifugated to recover and washed with PBS. Finally, cells were incubated with RNase for 30 min at 37 °C and stained with PI for 30 min at 4 °C. Flow cytometry (Beckman, CytoFLEX) and software ModFit LT 5.0 were utilized to perform the cell cycle analysis.

## QUANTIFICATION AND STATISTICAL ANALYSIS

### Statistics analysis

Statistical analyses were performed using GraphPad Prism Software (GraphPad Prism 8.0.1). Statistical analyses in mice were analyzed using the Mann-Whitney test. *P* values for mice survival were calculated using the Log-rank test. All other experiments were analyzed using unpaired, two-tailed Student’s t test, one-way or two-way ANOVA test. Error bars indicate ± SD. Statistical significance is denoted in figures by asterisks. \**P* < 0.05; \*\**P* < 0.01; \*\*\**P* < 0.001.).

---

**Table S1. Table S1 Bacterial strains and plasmids used in this study.**

**Table S2. Table S2 Primers used in this study.**

## REFERENCES

1. Behar, S.M., and Briken, V. (2019). Apoptosis inhibition by intracellular bacteria and its consequence on host immunity. Curr Opin Immunol 60, 103–110. 10.1016/j.coi.2019.05.007.

2. Dashzeveg, N., and Yoshida, K. (2015). Cell death decision by p53 via control of the mitochondrial membrane. Cancer Lett 367, 108–112. 10.1016/j.canlet.2015.07.019.

3. Jorgensen, I., Rayamajhi, M., and Miao, E.A. (2017). Programmed cell death as a defence against infection. Nat Rev Immunol 17, 151–164. 10.1038/nri.2016.147.

4. Singh, R., Letai, A., and Sarosiek, K. (2019). Regulation of apoptosis in health and disease: the balancing act of BCL-2 family proteins. Nat Rev Mol Cell Biol 20, 175–193. 10.1038/s41580-018-0089-8.

5. D’Arcy, M.S. (2019). Cell death: a review of the major forms of apoptosis, necrosis and autophagy. Cell Biol Int 43, 582–592. 10.1002/cbin.11137.

6. van Loo, G., and Bertrand, M.J.M. (2023). Death by TNF: a road to inflammation. Nat Rev Immunol 23, 289–303. 10.1038/s41577-022-00792-3.

7. Ai, Y., Meng, Y., Yan, B., Zhou, Q., and Wang, X. (2024). The biochemical pathways of apoptotic, necroptotic, pyroptotic, and ferroptotic cell death. Mol Cell 84, 170–179. 10.1016/j.molcel.2023.11.040.

8. Mao, W., Wang, Z., Wen, S., Lin, Y., Gu, J., Sun, J., Wang, H., Cao, Q., Xu, Y., Xu, X., and Cai, X. (2023). LRRC8A promotes Glaesserella parasuis cytolethal distending toxin-induced p53-dependent apoptosis in NPTr cells. Virulence 14, 2287339. 10.1080/21505594.2023.2287339.

9. Chumduri, C., Gurumurthy, R.K., Zietlow, R., and Meyer, T.F. (2016). Subversion of host genome integrity by bacterial pathogens. Nat Rev Mol Cell Biol 17, 659–673. 10.1038/nrm.2016.100.

10. Roos, W.P., and Kaina, B. (2006). DNA damage-induced cell death by apoptosis. Trends Mol Med 12, 440–450. 10.1016/j.molmed.2006.07.007.

11. Weinrauch, Y., and Zychlinsky, A. (1999). The induction of apoptosis by bacterial pathogens. Annu Rev Microbiol 53, 155–187. 10.1146/annurev.micro.53.1.155.

12. Martin, O.C.B., and Frisan, T. (2020). Bacterial Genotoxin-Induced DNA Damage and Modulation of the Host Immune Microenvironment. Toxins (Basel) 12, 63. 10.3390/toxins12020063.

13. Dziubanska-Kusibab, P.J., Berger, H., Battistini, F., Bouwman, B.A.M., Iftekhar, A., Katainen, R., Cajuso, T., Crosetto, N., Orozco, M., Aaltonen, L.A., and Meyer, T.F. (2020). Colibactin DNA-damage signature indicates mutational impact in colorectal cancer. Nat Med 26, 1063–1069. 10.1038/s41591-020-0908-2.

14. Jinadasa, R.N., Bloom, S.E., Weiss, R.S., and Duhamel, G.E. (2011). Cytolethal distending toxin: a conserved bacterial genotoxin that blocks cell cycle progression, leading to apoptosis of a broad range of mammalian cell lineages. Microbiology (Reading) 157, 1851–1875. 10.1099/mic.0.049536-0.

15. Cao, Y., Oh, J., Xue, M., Huh, W.J., Wang, J., Gonzalez-Hernandez, J.A., Rice, T.A., Martin, A.L., Song, D., Crawford, J.M., et al. (2022). Commensal microbiota from patients with inflammatory bowel disease produce genotoxic metabolites. Science 378, eabm3233. 10.1126/science.abm3233.

16. Dougherty, M.W., and Jobin, C. (2021). Shining a Light on Colibactin Biology. Toxins (Basel) 13. 10.3390/toxins13050346.

17. Bezine, E., Vignard, J., and Mirey, G. (2014). The cytolethal distending toxin effects on Mammalian cells: a DNA damage perspective. Cells 3, 592–615. 10.3390/cells3020592.

18. Wu, J., Sun, L., Chen, X., Du, F., Shi, H., Chen, C., and Chen, Z.J. (2013). Cyclic GMP-AMP is an endogenous second messenger in innate immune signaling by cytosolic DNA. Science 339, 826–830. 10.1126/science.1229963.

19. Mackenzie, K.J., Carroll, P., Martin, C.A., Murina, O., Fluteau, A., Simpson, D.J., Olova, N., Sutcliffe, H., Rainger, J.K., Leitch, A., et al. (2017). cGAS surveillance of micronuclei links genome instability to innate immunity. Nature 548, 461–465. 10.1038/nature23449.

20. Sze, A., Belgnaoui, S.M., Olagnier, D., Lin, R., Hiscott, J., and van Grevenynghe, J. (2013). Host restriction factor SAMHD1 limits human T cell leukemia virus type 1 infection of monocytes via STING-mediated apoptosis. Cell Host Microbe 14, 422–434. 10.1016/j.chom.2013.09.009.

21. Schock, S.N., Chandra, N.V., Sun, Y., Irie, T., Kitagawa, Y., Gotoh, B., Coscoy, L., and Winoto, A. (2017). Induction of necroptotic cell death by viral activation of the RIG-I or STING pathway. Cell Death Differ 24, 615–625. 10.1038/cdd.2016.153.

22. Reinert, L.S., Rashidi, A.S., Tran, D.N., Katzilieris-Petras, G., Hvidt, A.K., Gohr, M., Fruhwurth, S., Bodda, C., Thomsen, M.K., Vendelbo, M.H., et al. (2021). Brain immune cells undergo cGAS/STING-dependent apoptosis during herpes simplex virus type 1 infection to limit type I IFN production. J Clin Invest 131. 10.1172/JCI136824.

23. Gulen, M.F., Koch, U., Haag, S.M., Schuler, F., Apetoh, L., Villunger, A., Radtke, F., and Ablasser, A. (2017). Signalling strength determines proapoptotic functions of STING. Nat Commun 8, 427. 10.1038/s41467-017-00573-w.

24. Gaidt, M.M., Ebert, T.S., Chauhan, D., Ramshorn, K., Pinci, F., Zuber, S., O’Duill, F., Schmid-Burgk, J.L., Hoss, F., Buhmann, R., et al. (2017). The DNA Inflammasome in Human Myeloid Cells Is Initiated by a STING-Cell Death Program Upstream of NLRP3. Cell 171, 1110–1124 e1118. 10.1016/j.cell.2017.09.039.

25. Paludan, S.R., Reinert, L.S., and Hornung, V. (2019). DNA-stimulated cell death: implications for host defence, inflammatory diseases and cancer. Nat Rev Immunol 19, 141–153. 10.1038/s41577-018-0117-0.

26. Zheng, W., Liu, A., Xia, N., Chen, N., Meurens, F., and Zhu, J. (2023). How the Innate Immune DNA Sensing cGAS-STING Pathway Is Involved in Apoptosis. Int J Mol Sci 24. 10.3390/ijms24033029.

27. Prabakaran, T., Troldborg, A., Kumpunya, S., Alee, I., Marinkovic, E., Windross, S.J., Nandakumar, R., Narita, R., Zhang, B.C., Carstensen, M., et al. (2021). A STING antagonist modulating the interaction with STIM1 blocks ER-to-Golgi trafficking and inhibits lupus pathology. EBioMedicine 66, 103314. 10.1016/j.ebiom.2021.103314.

28. Dedoni, S., Olianas, M.C., and Onali, P. (2010). Interferon-beta induces apoptosis in human SH-SY5Y neuroblastoma cells through activation of JAK-STAT signaling and down-regulation of PI3K/Akt pathway. J Neurochem 115, 1421–1433. 10.1111/j.1471-4159.2010.07046.x.

29. Gao, W., Gao, J., Chen, L., Ren, Y., and Ma, J. (2019). Targeting XIST induced apoptosis of human osteosarcoma cells by activation of NF-kB/PUMA signal. Bioengineered 10, 261–270. 10.1080/21655979.2019.1631104.

30. van der Horst, D., Kurmasheva, N., Marqvorsen, M.H.S., Assil, S., Rahimic, A.H.F., Kollmann, C.F., Silva da Costa, L., Wu, Q., Zhao, J., Cesari, E., et al. (2024). SAM68 directs STING signaling to apoptosis in macrophages. Commun Biol 7, 283. 10.1038/s42003-024-05969-1.

31. Gao, D., Li, T., Li, X.D., Chen, X., Li, Q.Z., Wight-Carter, M., and Chen, Z.J. (2015). Activation of cyclic GMP-AMP synthase by self-DNA causes autoimmune diseases. Proc Natl Acad Sci U S A 112, E5699–5705. 10.1073/pnas.1516465112.

32. Li, C., Liu, W., Wang, F., Hayashi, T., Mizuno, K., Hattori, S., Fujisaki, H., and Ikejima, T. (2021). DNA damage-triggered activation of cGAS-STING pathway induces apoptosis in human keratinocyte HaCaT cells. Mol Immunol 131, 180–190. 10.1016/j.molimm.2020.12.037.

33. Creagh, E.M., Conroy, H., and Martin, S.J. (2003). Caspase-activation pathways in apoptosis and immunity. Immunol Rev 193, 10–21. 10.1034/j.1600-065x.2003.00048.x.

34. Karaba, S.M., White, R.C., and Cianciotto, N.P. (2013). Stenotrophomonas maltophilia encodes a type II protein secretion system that promotes detrimental effects on lung epithelial cells. Infect Immun 81, 3210–3219. 10.1128/IAI.00546-13.

35. Nas, M.Y., White, R.C., DuMont, A.L., Lopez, A.E., and Cianciotto, N.P. (2019). Stenotrophomonas maltophilia Encodes a VirB/VirD4 Type IV Secretion System That Modulates Apoptosis in Human Cells and Promotes Competition against Heterologous Bacteria, Including Pseudomonas aeruginosa. Infect Immun 87. 10.1128/IAI.00457-19.

36. Samba-Louaka, A., Nougayrede, J.P., Watrin, C., Oswald, E., and Taieb, F. (2009). The enteropathogenic Escherichia coli effector Cif induces delayed apoptosis in epithelial cells. Infect Immun 77, 5471–5477. 10.1128/IAI.00860-09.

37. Nougayrede, J.P., and Donnenberg, M.S. (2004). Enteropathogenic Escherichia coli EspF is targeted to mitochondria and is required to initiate the mitochondrial death pathway. Cell Microbiol 6, 1097–1111. 10.1111/j.1462-5822.2004.00421.x.

38. Banga, S., Gao, P., Shen, X., Fiscus, V., Zong, W.X., Chen, L., and Luo, Z.Q. (2007). Legionella pneumophila inhibits macrophage apoptosis by targeting pro-death members of the Bcl2 protein family. Proc Natl Acad Sci U S A 104, 5121–5126. 10.1073/pnas.0611030104.

39. Suarez, G., Sierra, J.C., Sha, J., Wang, S., Erova, T.E., Fadl, A.A., Foltz, S.M., Horneman, A.J., and Chopra, A.K. (2008). Molecular characterization of a functional type VI secretion system from a clinical isolate of Aeromonas hydrophila. Microb Pathog 44, 344–361. 10.1016/j.micpath.2007.10.005.

40. Zhou, Y., Tao, J., Yu, H., Ni, J., Zeng, L., Teng, Q., Kim, K.S., Zhao, G.P., Guo, X., and Yao, Y. (2012). Hcp family proteins secreted via the type VI secretion system coordinately regulate Escherichia coli K1 interaction with human brain microvascular endothelial cells. Infect Immun 80, 1243–1251. 10.1128/IAI.05994-11.

41. Pukatzki, S., Ma, A.T., Sturtevant, D., Krastins, B., Sarracino, D., Nelson, W.C., Heidelberg, J.F., and Mekalanos, J.J. (2006). Identification of a conserved bacterial protein secretion system in Vibrio cholerae using the Dictyostelium host model system. Proc Natl Acad Sci U S A 103, 1528–1533. 10.1073/pnas.0510322103.

42. Cascales, E., and Cambillau, C. (2012). Structural biology of type VI secretion systems. Philos Trans R Soc Lond B Biol Sci 367, 1102–1111. 10.1098/rstb.2011.0209.

43. Ma, A.T., and Mekalanos, J.J. (2010). In vivo actin cross-linking induced by Vibrio cholerae type VI secretion system is associated with intestinal inflammation. Proc Natl Acad Sci U S A 107, 4365–4370. 10.1073/pnas.0915156107.

44. Monjaras Feria, J., and Valvano, M.A. (2020). An Overview of Anti-Eukaryotic T6SS Effectors. Front Cell Infect Microbiol 10, 584751. 10.3389/fcimb.2020.584751.

45. Wan, B., Zhang, Q., Ni, J., Li, S., Wen, D., Li, J., Xiao, H., He, P., Ou, H.Y., Tao, J., et al. (2017). Type VI secretion system contributes to Enterohemorrhagic Escherichia coli virulence by secreting catalase against host reactive oxygen species (ROS). PLoS Pathog 13, e1006246. 10.1371/journal.ppat.1006246.

46. Zong, B., Zhang, Y., Wang, X., Liu, M., Zhang, T., Zhu, Y., Zheng, Y., Hu, L., Li, P., Chen, H., and Tan, C. (2019). Characterization of multiple type-VI secretion system (T6SS) VgrG proteins in the pathogenicity and antibacterial activity of porcine extra-intestinal pathogenic Escherichia coli. Virulence 10, 118–132. 10.1080/21505594.2019.1573491.

47. Whitney, J.C., Chou, S., Russell, A.B., Biboy, J., Gardiner, T.E., Ferrin, M.A., Brittnacher, M., Vollmer, W., and Mougous, J.D. (2013). Identification, structure, and function of a novel type VI secretion peptidoglycan glycoside hydrolase effector-immunity pair. J Biol Chem 288, 26616–26624. 10.1074/jbc.M113.488320.

48. Russell, A.B., Hood, R.D., Bui, N.K., LeRoux, M., Vollmer, W., and Mougous, J.D. (2011). Type VI secretion delivers bacteriolytic effectors to target cells. Nature 475, 343–347. 10.1038/nature10244.

49. Russell, A.B., LeRoux, M., Hathazi, K., Agnello, D.M., Ishikawa, T., Wiggins, P.A., Wai, S.N., and Mougous, J.D. (2013). Diverse type VI secretion phospholipases are functionally plastic antibacterial effectors. Nature 496, 508–512. 10.1038/nature12074.

50. Miyata, S.T., Kitaoka, M., Brooks, T.M., McAuley, S.B., and Pukatzki, S. (2011). Vibrio cholerae requires the type VI secretion system virulence factor VasX to kill Dictyostelium discoideum. Infect Immun 79, 2941–2949. 10.1128/IAI.01266-10.

51. Jana, B., Fridman, C.M., Bosis, E., and Salomon, D. (2019). A modular effector with a DNase domain and a marker for T6SS substrates. Nat Commun 10, 3595. 10.1038/s41467-019-11546-6.

52. Pissaridou, P., Allsopp, L.P., Wettstadt, S., Howard, S.A., Mavridou, D.A.I., and Filloux, A. (2018). The Pseudomonas aeruginosa T6SS-VgrG1b spike is topped by a PAAR protein eliciting DNA damage to bacterial competitors. Proc Natl Acad Sci U S A 115, 12519–12524. 10.1073/pnas.1814181115.

53. Song, L., Pan, J., Yang, Y., Zhang, Z., Cui, R., Jia, S., Wang, Z., Yang, C., Xu, L., Dong, T.G., et al. (2021). Contact-independent killing mediated by a T6SS effector with intrinsic cell-entry properties. Nat Commun 12, 423. 10.1038/s41467-020-20726-8.

54. Song, L., Xu, L., Wu, T., Shi, Z., Kareem, H.A., Wang, Z., Dai, Q., Guo, C., Pan, J., Yang, M., et al. (2024). Trojan horselike T6SS effector TepC mediates both interference competition and exploitative competition. ISME J 18. 10.1093/ismejo/wrad028.

55. Jiang, F., Wang, X., Wang, B., Chen, L., Zhao, Z., Waterfield, N.R., Yang, G., and Jin, Q. (2016). The Pseudomonas aeruginosa Type VI Secretion PGAP1-like Effector Induces Host Autophagy by Activating Endoplasmic Reticulum Stress. Cell Rep 16, 1502–1509. 10.1016/j.celrep.2016.07.012.

56. Jiang, F., Waterfield, N.R., Yang, J., Yang, G., and Jin, Q. (2014). A Pseudomonas aeruginosa type VI secretion phospholipase D effector targets both prokaryotic and eukaryotic cells. Cell Host Microbe 15, 600–610. 10.1016/j.chom.2014.04.010.

57. Ma, L.S., Hachani, A., Lin, J.S., Filloux, A., and Lai, E.M. (2014). Agrobacterium tumefaciens deploys a superfamily of type VI secretion DNase effectors as weapons for interbacterial competition in planta. Cell Host Microbe 16, 94–104. 10.1016/j.chom.2014.06.002.

58. Luo, J., Chu, X., Jie, J., Sun, Y., Guan, Q., Li, D., Luo, Z.Q., and Song, L. (2023). Acinetobacter baumannii Kills Fungi via a Type VI DNase Effector. mBio 14, e0342022. 10.1128/mbio.03420-22.

59. Kleinstiver, B.P., Wolfs, J.M., Kolaczyk, T., Roberts, A.K., Hu, S.X., and Edgell, D.R. (2012). Monomeric site-specific nucleases for genome editing. Proc Natl Acad Sci U S A 109, 8061–8066. 10.1073/pnas.1117984109.

60. Ma, A.T., McAuley, S., Pukatzki, S., and Mekalanos, J.J. (2009). Translocation of a Vibrio cholerae type VI secretion effector requires bacterial endocytosis by host cells. Cell Host Microbe 5, 234–243. 10.1016/j.chom.2009.02.005.

61. Rogakou, E.P., Pilch, D.R., Orr, A.H., Ivanova, V.S., and Bonner, W.M. (1998). DNA double-stranded breaks induce histone H2AX phosphorylation on serine 139. J Biol Chem 273, 5858–5868. 10.1074/jbc.273.10.5858.

62. Zierhut, C., and Funabiki, H. (2020). Regulation and Consequences of cGAS Activation by Self-DNA. Trends Cell Biol 30, 594–605. 10.1016/j.tcb.2020.05.006.

63. Liu, H., Zhang, H., Wu, X., Ma, D., Wu, J., Wang, L., Jiang, Y., Fei, Y., Zhu, C., Tan, R., et al. (2018). Nuclear cGAS suppresses DNA repair and promotes tumorigenesis. Nature 563, 131–136. 10.1038/s41586-018-0629-6.

64. Newton, K., Dixit, V.M., and Kayagaki, N. (2021). Dying cells fan the flames of inflammation. Science 374, 1076–1080. 10.1126/science.abi5934.

65. Zhang, Y., and Bliska, J.B. (2005). Role of macrophage apoptosis in the pathogenesis of Yersinia. Curr Top Microbiol Immunol 289, 151–173. 10.1007/3-540-27320-4_7.

66. Hering, N.A., Richter, J.F., Krug, S.M., Gunzel, D., Fromm, A., Bohn, E., Rosenthal, R., Bucker, R., Fromm, M., Troeger, H., and Schulzke, J.D. (2011). Yersinia enterocolitica induces epithelial barrier dysfunction through regional tight junction changes in colonic HT-29/B6 cell monolayers. Lab Invest 91, 310–324. 10.1038/labinvest.2010.180.

67. Wang, D., Zhao, H., Shen, Y., and Chen, Q. (2022). A Variety of Nucleic Acid Species Are Sensed by cGAS, Implications for Its Diverse Functions. Front Immunol 13, 826880. 10.3389/fimmu.2022.826880.

68. Sana, T.G., Flaugnatti, N., Lugo, K.A., Lam, L.H., Jacobson, A., Baylot, V., Durand, E., Journet, L., Cascales, E., and Monack, D.M. (2016). Salmonella Typhimurium utilizes a T6SS-mediated antibacterial weapon to establish in the host gut. Proc Natl Acad Sci U S A 113, E5044–5051. 10.1073/pnas.1608858113.

69. Zhao, W., Caro, F., Robins, W., and Mekalanos, J.J. (2018). Antagonism toward the intestinal microbiota and its effect on Vibrio cholerae virulence. Science 359, 210–213. 10.1126/science.aap8775.

70. Lin, J., Xu, L., Yang, J., Wang, Z., and Shen, X. (2021). Beyond dueling: roles of the type VI secretion system in microbiome modulation, pathogenesis and stress resistance. Stress Biol 1, 11. 10.1007/s44154-021-00008-z.

71. Zhang, G., Wang, J., Zhao, Z., Xin, T., Fan, X., Shen, Q., Raheem, A., Lee, C.R., Jiang, H., and Ding, J. (2022). Regulated necrosis, a proinflammatory cell death, potentially counteracts pathogenic infections. Cell Death Dis 13, 637. 10.1038/s41419-022-05066-3.

72. Raymond, B., Young, J.C., Pallett, M., Endres, R.G., Clements, A., and Frankel, G. (2013). Subversion of trafficking, apoptosis, and innate immunity by type III secretion system effectors. Trends Microbiol 21, 430–441. 10.1016/j.tim.2013.06.008.

73. Teng, Y.T., and Hu, W. (2003). Expression cloning of a periodontitis-associated apoptotic effector, cagE homologue, in Actinobacillus actinomycetemcomitans. Biochem Biophys Res Commun 303, 1086–1094. 10.1016/s0006-291x(03)00471-6.

74. DuMont, A.L., and Cianciotto, N.P. (2017). Stenotrophomonas maltophilia Serine Protease StmPr1 Induces Matrilysis, Anoikis, and Protease-Activated Receptor 2 Activation in Human Lung Epithelial Cells. Infect Immun 85. 10.1128/IAI.00544-17.

75. Orth, K., Palmer, L.E., Bao, Z.Q., Stewart, S., Rudolph, A.E., Bliska, J.B., and Dixon, J.E. (1999). Inhibition of the mitogen-activated protein kinase kinase superfamily by a Yersinia effector. Science 285, 1920–1923. 10.1126/science.285.5435.1920.

76. Hersh, D., Monack, D.M., Smith, M.R., Ghori, N., Falkow, S., and Zychlinsky, A. (1999). The Salmonella invasin SipB induces macrophage apoptosis by binding to caspase-1. Proc Natl Acad Sci U S A 96, 2396–2401. 10.1073/pnas.96.5.2396.

77. Senerovic, L., Tsunoda, S.P., Goosmann, C., Brinkmann, V., Zychlinsky, A., Meissner, F., and Kolbe, M. (2012). Spontaneous formation of IpaB ion channels in host cell membranes reveals how Shigella induces pyroptosis in macrophages. Cell Death Dis 3, e384. 10.1038/cddis.2012.124.

78. Shenker, B.J., Walker, L.M., Zekavat, A., Weiss, R.H., and Boesze-Battaglia, K. (2020). The Cell-Cycle Regulatory Protein p21CIP1/WAF1 Is Required for Cytolethal Distending Toxin (Cdt)-Induced Apoptosis. Pathogens 9, 38.

79. Ku, J.W.K., Chen, Y., Lim, B.J.W., Gasser, S., Crasta, K.C., and Gan, Y.H. (2020). Bacterial-induced cell fusion is a danger signal triggering cGAS-STING pathway via micronuclei formation. Proc Natl Acad Sci U S A 117, 15923–15934. 10.1073/pnas.2006908117.

80. Zhu, L., Xu, L., Wang, C., Li, C., Li, M., Liu, Q., Wang, X., Yang, W., Pan, D., Hu, L., et al. (2021). T6SS translocates a micropeptide to suppress STING-mediated innate immunity by sequestering manganese. Proc Natl Acad Sci U S A 118. 10.1073/pnas.2103526118.

81. Huyghe, J., Priem, D., and Bertrand, M.J.M. (2023). Cell death checkpoints in the TNF pathway. Trends Immunol 44, 628–643. 10.1016/j.it.2023.05.007.

82. Elinav, E., Nowarski, R., Thaiss, C.A., Hu, B., Jin, C., and Flavell, R.A. (2013). Inflammation-induced cancer: crosstalk between tumours, immune cells and microorganisms. Nat Rev Cancer 13, 759–771. 10.1038/nrc3611.

83. Xiang, Y., Zhang, C., Wang, J., Cheng, Y., Wang, L., Tong, Y., and Yan, D. (2023). Identification of host gene-microbiome associations in colorectal cancer patients using mendelian randomization. J Transl Med 21, 535. 10.1186/s12967-023-04335-9.

84. Kang, J., Sun, M., Chang, Y., Chen, H., Zhang, J., Liang, X., and Xiao, T. (2023). Butyrate ameliorates colorectal cancer through regulating intestinal microecological disorders. Anticancer Drugs 34, 227–237. 10.1097/CAD.0000000000001413.

85. Sun, T., Liu, S., Zhou, Y., Yao, Z., Zhang, D., Cao, S., Wei, Z., Tan, B., Li, Y., Lian, Z., and Wang, S. (2017). Evolutionary biologic changes of gut microbiota in an ’adenoma-carcinoma sequence’ mouse colorectal cancer model induced by 1, 2-Dimethylhydrazine. Oncotarget 8, 444–457. 10.18632/oncotarget.13443.

86. Singh, R., Chandrashekharappa, S., Bodduluri, S.R., Baby, B.V., Hegde, B., Kotla, N.G., Hiwale, A.A., Saiyed, T., Patel, P., Vijay-Kumar, M., et al. (2019). Enhancement of the gut barrier integrity by a microbial metabolite through the Nrf2 pathway. Nat Commun 10, 89. 10.1038/s41467-018-07859-7.

87. Zhang, X., Yu, D., Wu, D., Gao, X., Shao, F., Zhao, M., Wang, J., Ma, J., Wang, W., Qin, X., et al. (2023). Tissue-resident Lachnospiraceae family bacteria protect against colorectal carcinogenesis by promoting tumor immune surveillance. Cell Host Microbe 31, 418–432 e418. 10.1016/j.chom.2023.01.013.

88. Xu, L., Li, M., Yang, Y., Zhang, C., Xie, Z., Tang, J., Shi, Z., Chen, S., Li, G., Gu, Y., et al. (2022). Salmonella Induces the cGAS-STING-Dependent Type I Interferon Response in Murine Macrophages by Triggering mtDNA Release. mBio 13, e0363221. 10.1128/mbio.03632-21.

