## Supplemental Figs for "A T6SS DNase effector induces nuclear DNA Damage to trigger apoptosis via activation of the cGAS-STING-TNF axis"

### Supplementary Figures and Legends

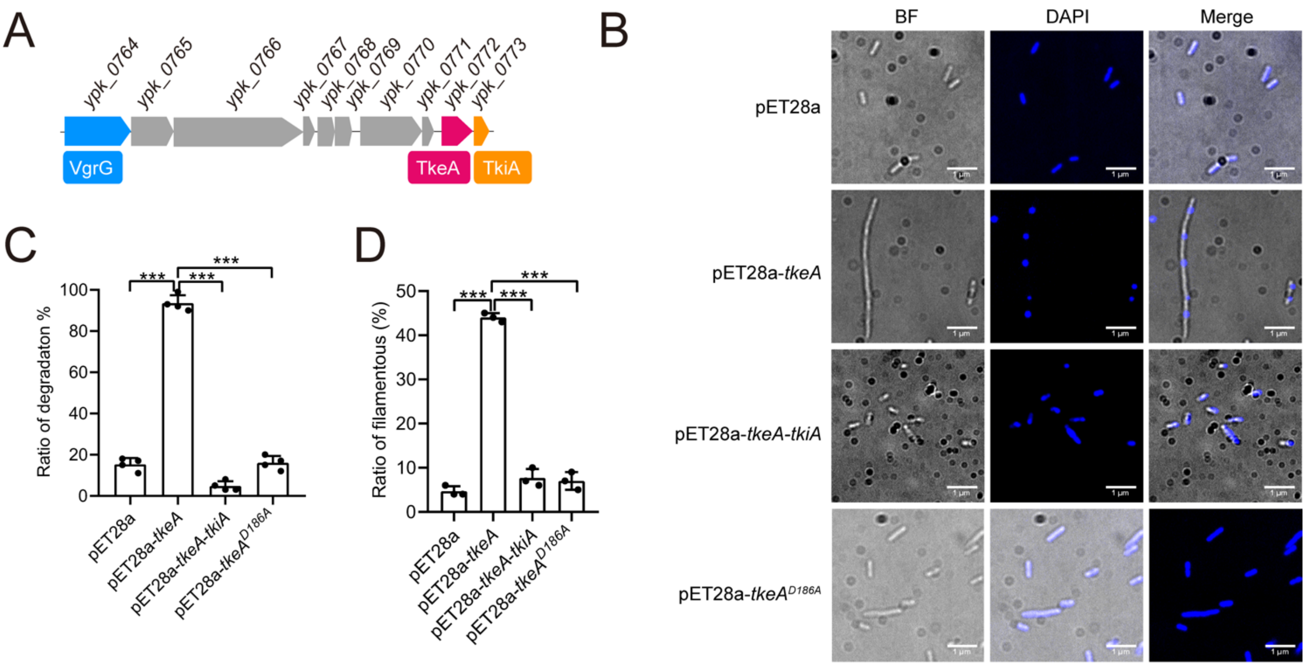


**Figure S1. TkeA degrades *E. coli* genome DNA. Related to Figure 1**

(A) Structure of the *Yptb* orphan gene cluster (From *ypk_0764* to *ypk_0773*). The *vgrG* gene (*ypk_0764*), *tkeA* gene (*ypk_0772*) and *tkiA* gene (*ypk_0773*) are colored in blue, yellow and red respectively.

(B) Detection of the loss of DNA staining (DAPI) in *E. coli* cells expression TkeA, TkeA$^{D186A}$ and co-expressing TkeA-TkiA at 4 h after IPTG induction. Fluorescence microscopy was performed to visualize the genome degradation. Scale bar, 1 µm.

(C) and (D) The quantification in (A) of the degradation and filamentous were calculated. Data are from 3 biological replicates.

*P* values calculated using one-way analysis of variance (ANOVA) for multiple comparisons.

Error bars represent ± SD. ****P* < 0.001.

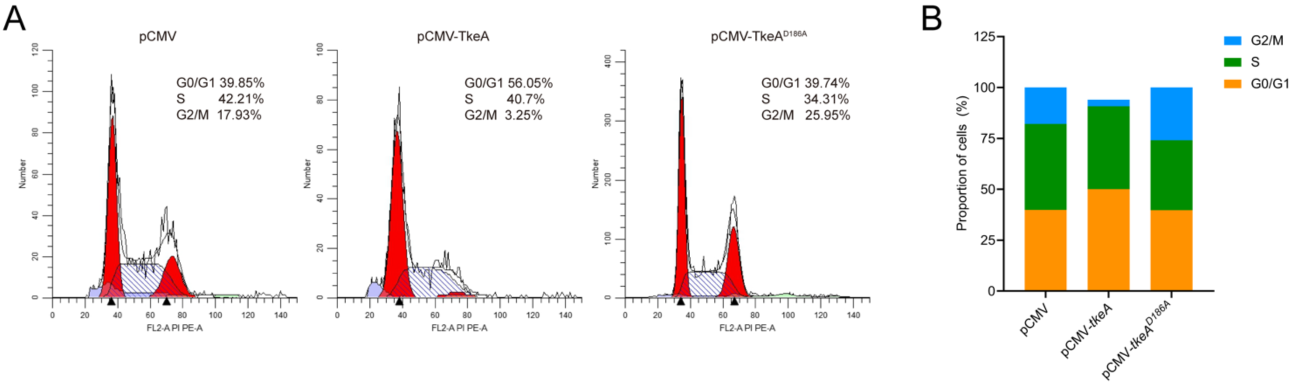


**Figure S2. TkeA induces cell cycle arrest at the G1 phase in HeLa cells. Related to Figure 2**

(A)HeLa cells were transfected with pCMV (Control), pCMV-*tkeA* vector and pCMV-*tkeA*$^{D186A}$ and were collected and stained with PI. The cell cycle was detected by flow cytometry.

(B) The quantification in (A) of the cell cycle.

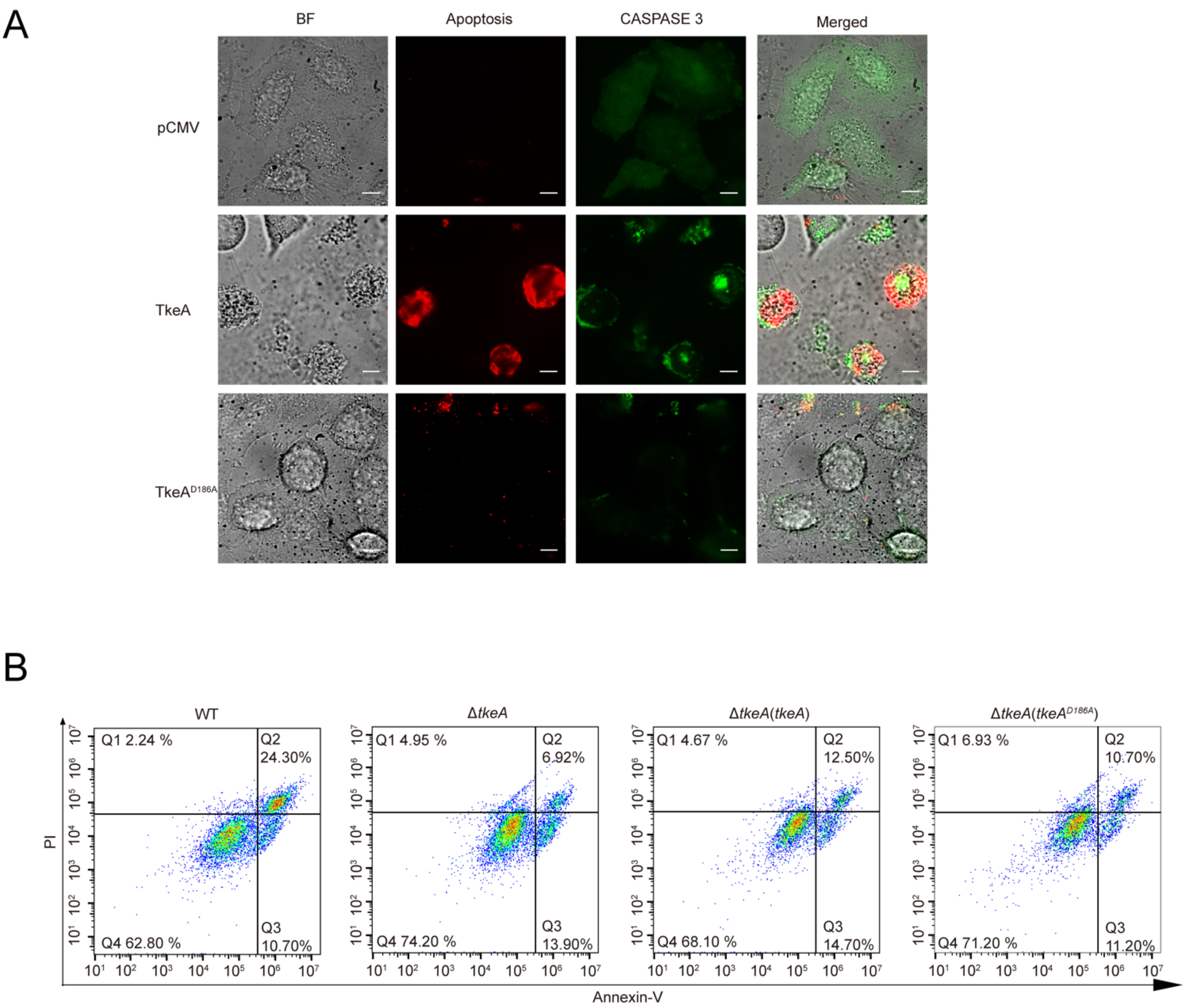


**Figure S3. TkeA activates CASPASE 3 and apoptosis in host cells. Related to Figure 3**

(A) CASPASE 3 activated and apoptosis test in HeLa cells. HeLa cells transfected with pCMV, pCMV-*tkeA* and pCMV-*tkeA*$^{D186A}$ for 24 h were stained with GreenNuc™ CASPASE 3 and Annexin V-mCherry. Images are representative cells from the same field of view. Fluorescence microscopy was performed to visualize the activation of CASPASE 3 and apoptosis. Scale bar, 500 µm.

(B) HeLa cells infected with the indicated *Yptb* strains were collected and stained with Annexin V/PI. Flow cytometry was used to identify the cell apoptosis.

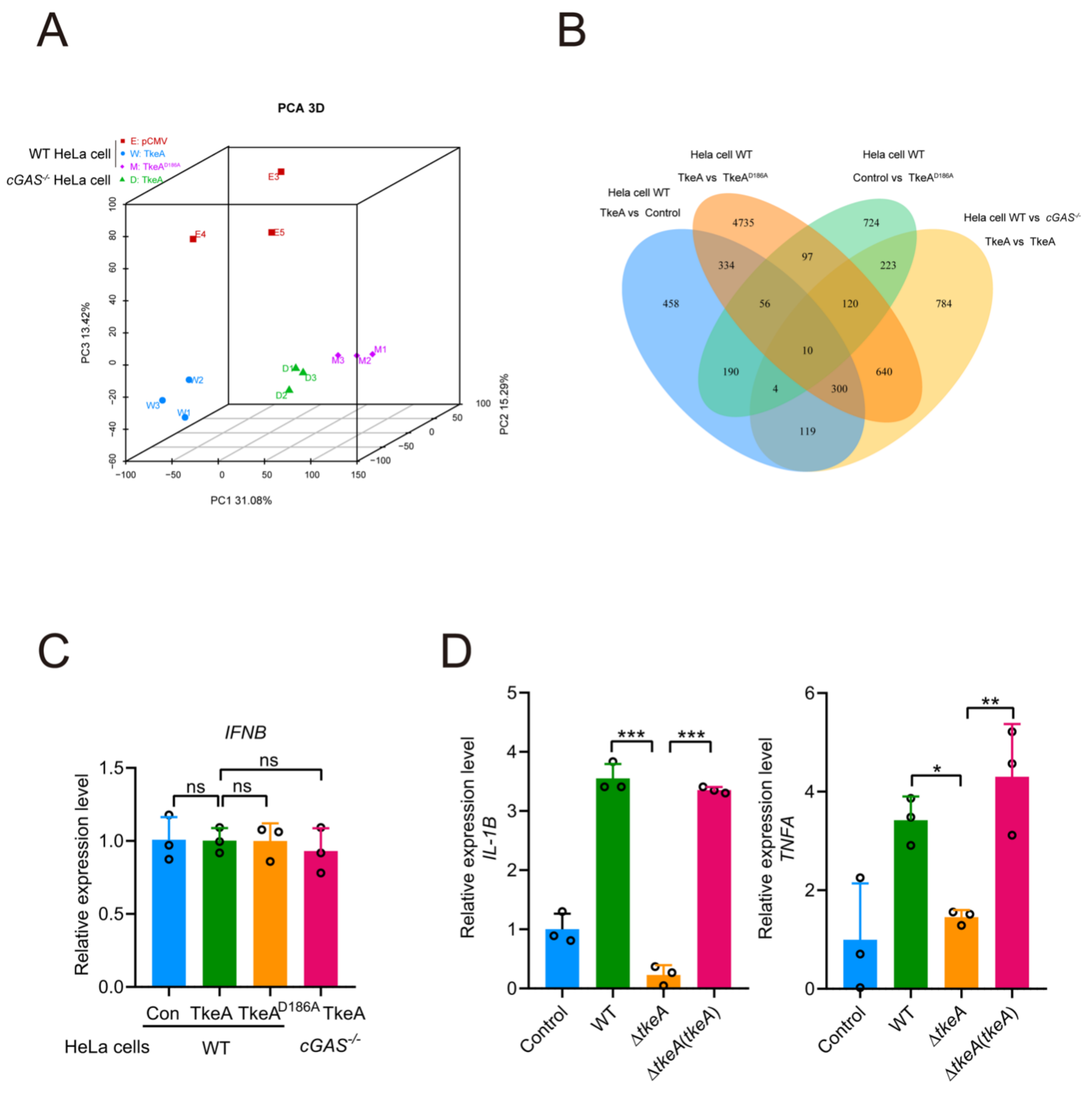


**Figure S4. The cGAS-STING-TNF signaling pathway is implicated in TkeA-induced apoptosis. Related to Figure 5**

(A) WT HeLa cells transfected with pCMV, pCMV-*tkeA* and pCMV-*tkeA*$^{D186A}$, and *cGAS*$^{-/-}$ HeLa cells transfected with pCMV-*tkeA*. RNA isolated from these cells was subject to RNA-seq. Principal component analysis (PCA) was used to compare the four groups. n = 3.

(B) A Venn diagram was used to illustrate the unions, intersections and distinctions among four groups. n = 3.

(C) qRT-PCR analysis of gene expression in WT HeLa cells transfected with pCMV, pCMV-*tkeA* and pCMV-*tkeA*$^{D186A}$, and cGAS$^{-/-}$HeLa cells transfected with pCMV-*tkeA*. n = 3.

(D) qRT-PCR analysis of gene expression in WT HeLa cells infected with *Yptb* WT, Δ*tkeA* and Δ*tkeA(tkeA)* for 4 h at an MOI of 100. n = 3.

*P* values calculated using one-way ANOVA for multiple comparisons.

Error bars represent ± SD. *$P < 0.05$; ***$P < 0.001$. ns, not significant.

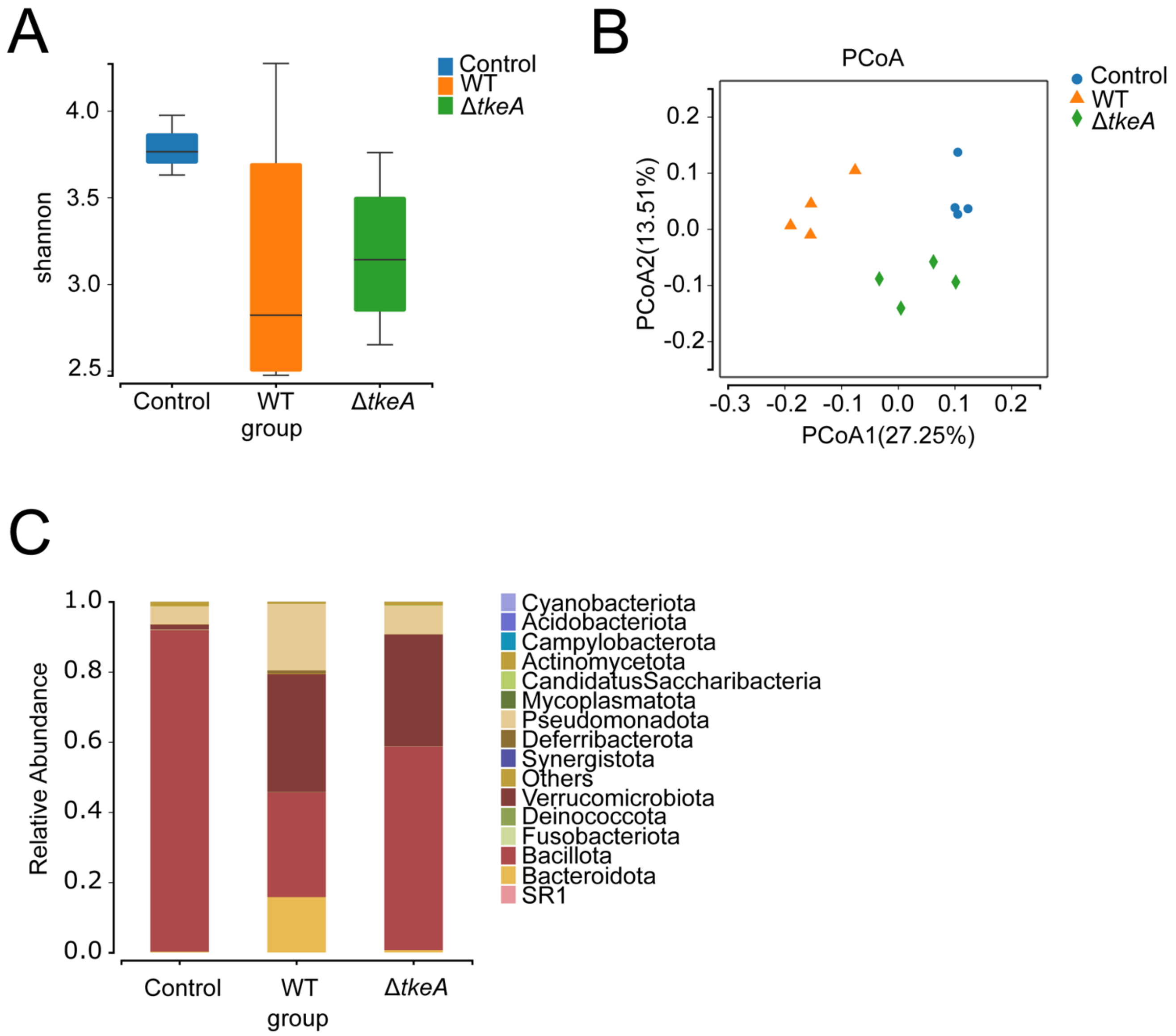


**Figure S5. The analysis of 16S rRNA gene amplicon of the microbiota of mice infected with different *Yptb* strains. Related to Figure 6**

(A) mice were orally gavaged with $10^9$ CFUs of different *Yptb* strains. Alpha diversity of the gut microbiota with the Shannon index in the three groups. The horizontal bars within boxes represent medians.

(B) CPCoA with Bray-Curtis distance showing the beta diversity ($P$<0.001, PERMANOVA by Adonis).

(C) Phylum-level distribution of the native gut microbiota in three groups.

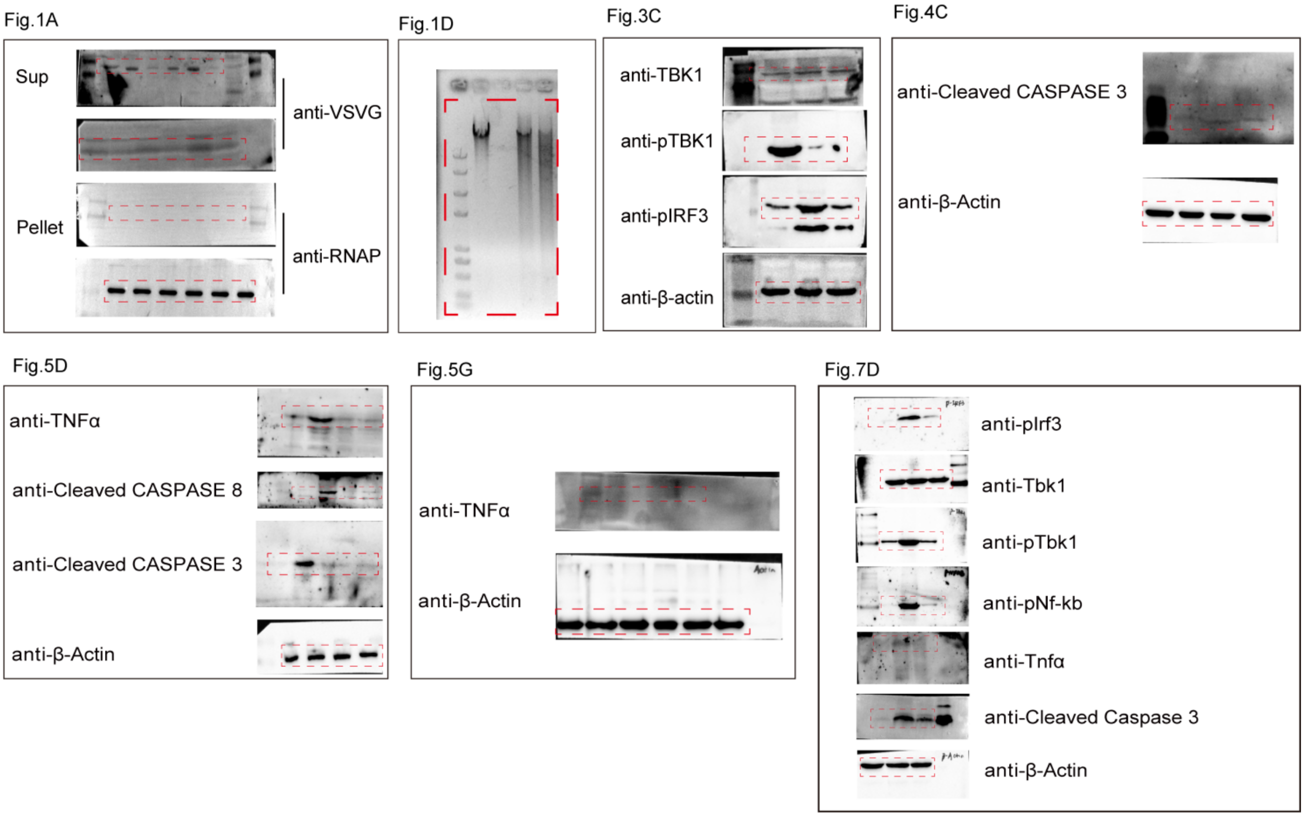


**Figure S6. Uncropped versions of immunoblotting results. Related to Figure 1, 3, 4, 5 and 7**
